# Single-cell transcriptomics reveals a conserved metaplasia program in pancreatic injury

**DOI:** 10.1101/2021.04.09.439243

**Authors:** Zhibo Ma, Nikki K. Lytle, Bob Chen, Nidhi Jyotsana, Sammy Weiser Novak, Charles J. Cho, Leah Caplan, Olivia Ben-Levy, Abigail C. Neininger, Dylan T. Burnette, Vincent Q. Trinh, Marcus C.B. Tan, Emilee A. Patterson, Rafael Arrojo e Drigo, Rajshekhar R. Giraddi, Cynthia Ramos, Anna L. Means, Ichiro Matsumoto, Uri Manor, Jason C. Mills, James R. Goldenring, Ken S. Lau, Geoffrey M. Wahl, Kathleen E. DelGiorno

**Author notes:** Correspondence to: Kathleen DelGiorno. Pfizer Inc., San Diego, CA, 92121. **Biorxiv doi**: https://doi.org/10.1101/2021.04.09.439243. **Author contributions**: Conceptualization: K.E.D. and Z.M. Formal analysis: Z.M, N.K.L., B.C., N.J., S.W.N., C.J.C., D.T.B., V.Q.T., M.T., R.A., R.R.G., K.E.D. Funding acquisition: M.T., U.M., K.S.L, G.M.W, K.E.D. Investigation: Z.M., N.K.L., B.C., N.J., S.W.N., C.J.C., A.C.N., E.A.P, C.R., A.L.M., J.R.G. Project administration: K.E.D. Resources: J.C.M., K.S.L., I.M. Software: Z.M., B.C., S.W.N. Supervision: D.T.B., M.T., R.A., U.M., J.C.M., K.S.L, G.M.W., K.E.D Visualization: Z.M., N.K.L., B.C., S.W.N., C.J.C., A.C.N., D.T.B., R.A., K.E.D. **Data and materials availability**: Sequencing data that support this study is available in the GEO archive (GSE172380).

## Abstract

**BACKGROUND & AIMS:** Acinar to ductal metaplasia (ADM) occurs in the pancreas in response to tissue injury and is a potential precursor for adenocarcinoma. The goal of these studies was to define the populations arising from ADM, the associated transcriptional changes, and markers of disease progression.

**METHODS:** Acinar cells were lineage-traced with enhanced yellow fluorescent protein (EYFP) to follow their fate upon injury. Transcripts of over 13,000 EYFP+ cells were determined using single-cell RNA sequencing (scRNA-seq). Developmental trajectories were generated. Data were compared to gastric metaplasia, *Kras*^*G12D*^-induced neoplasia, and human pancreatitis. Results were confirmed by immunostaining and electron microscopy. *Kras*^*G12D*^ was expressed in injury-induced ADM using several inducible Cre drivers. Surgical specimens of chronic pancreatitis from 15 patients were evaluated by immunostaining.

**RESULTS:** scRNA-seq of ADM revealed emergence of a mucin/ductal population resembling gastric pyloric metaplasia. Lineage trajectories suggest that some pyloric metaplasia cells can generate tuft and enteroendocrine cells (EECs). Comparison to *Kras*^*G12D*^-induced ADM identifies populations associated with disease progression. Activation of *Kras*^*G12D*^ expression in HNF1B+ or POU2F3+ ADM populations leads to neoplastic transformation and formation of MUC5AC+ gastric-pit-like cells. Human pancreatitis samples also harbor pyloric metaplasia with a similar transcriptional phenotype.

**CONCLUSIONS:** Under conditions of chronic injury, acinar cells undergo a pyloric-type metaplasia to mucinous progenitor-like populations, which seed disparate tuft cell and EEC lineages. ADM-derived EEC subtypes are diverse. *Kras*^*G12D*^ expression is sufficient to drive neoplasia from injury-induced ADM and offers an alternative origin for tumorigenesis. This program is conserved in human pancreatitis, providing insight into early events in pancreas diseases.

## INTRODUCTION

Damage to the exocrine compartment of the pancreas results in infiltration of immune cells, stromal deposition, loss of acinar cells, and the emergence of a metaplastic cell lineage pathologically defined by the aberrant appearance of ducts termed ‘acinar to ductal metaplasia’ (ADM)^1^. ADM is a reparative program in which digestive enzyme-producing acinar cells transdifferentiate to cells with ductal organization as part of a critical acute phase program (paligenosis) that enables tissue reconstruction following injury^2, 3^.

Under conditions of sustained injury, inherited mutation, environmental factors, or idiopathic stimuli, patients can develop chronic pancreatitis, a known risk factor for pancreatic ductal adenocarcinoma (PDAC). ADM is a pathognomonic epithelial change during chronic pancreatitis and occurs in response to oncogenic mutation in the exocrine tissue, identifying it as a pre-neoplastic event thought to be critical for eventual malignant transformation^1, 4^. The induction of injury and inflammation-induced ADM, therefore, may be one factor by which chronic pancreatitis increases the risk of PDAC.

We have recently shown that ADM results in the transdifferentiation of acinar cells to tuft cells, which are rare chemosensory cells typically absent from the normal mouse pancreas^5^. Using a combination of low-input RNA sequencing strategies, super resolution microscopy, and genetically engineered mouse models, we demonstrated that tuft cells inhibit pancreatic tumorigenesis by modulating tissue stroma^5-7^. These data demonstrate that rare ADM-derived cell types can significantly impact disease progression. Given the role of ADM in disease initiation and the potentially critical link between injury and tumorigenesis, we combined lineage tracing and single-cell RNA sequencing (scRNA-seq) to identify populations arising from ADM.

Using this strategy, we identified the emergence of tuft cells, enteroendocrine cells, and mucin/ductal cell populations bearing gene signatures similar to spasmolytic polypeptide-expressing metaplasia (SPEM) in the stomach *(Tff2+Muc6+Gif+)*. Gene expression analysis of these populations in combination with trajectory inference and RNA velocity modeling suggests varying proclivity for seeding disparate differentiated tuft and enteroendocrine cell lineages. Formation of distinct cell types was confirmed by marker immunostaining and electron microscopy for cell type-specific structures. Our data were compared to *Kras*^*G12D*^-induced ADM and *Kras*^*G12D*^ expression was activated in injury-induced ADM populations using several inducible Cre drivers. Finally, we compared our dataset to published single nucleus RNA sequencing (sNuc-seq) data from human pancreatitis and identified conserved processes^8^. These studies identify the emergence of previously undescribed acinar-derived cell types in pancreatic injury, metaplasia transcriptional programs shared between organ systems, population changes between injury and tumorigenesis, and a conserved response to exocrine injury in human disease.

## RESULTS

### Lineage tracing and scRNA-seq identifies epithelial heterogeneity in injury-induced ADM

The goal of these studies was to evaluate cellular heterogeneity in ADM resulting from chronic injury in genetically wild type pancreatitis. To this end, we conducted lineage tracing by labeling adult pancreatic acinar cells with EYFP in *Ptf1a*^*CreERTM*^*;Rosa*^*LSL-YFP/+*^ (CERTY) mice. We then induced pancreatic injury with repeated injection of the cholecystokinin ortholog caerulein; resulting in metaplasia, cell death, and inflammation^5, 9^. Mice were sacrificed after either 2 or 4 weeks of treatment and pancreata were collected (Figure 1A, S1A). We isolated EYFP+ cells by fluorescence activated cell sorting (FACS) and profiled the transcriptome with 10x Genomics 3’ scRNA sequencing (Figure 1A, S1B). We used lineage tracing to identify metaplastic populations derived from acinar cells (i.e., ADM populations), but did not perform additional selections in order to agnostically profile the EYFP-labeled population. The combined libraries from the 2- and 4-week time points (2 mice per time point) contained a total of 21,140 cells with an average of 2,782 genes and 17,253 unique molecular identifiers detected per cell (File S1). Clusters were annotated by examining both classic cell type markers and previously published single-cell gene signatures in pancreas tissue^10^. Cells clustered mainly by cell type and data from the two time-points were found to largely overlap (Figure S1C). We identified both epithelial and stromal populations in our dataset, however EYFP transcripts were found to be specific to Krt8+ epithelial populations (Figure S2A-B).

**Figure 1.**
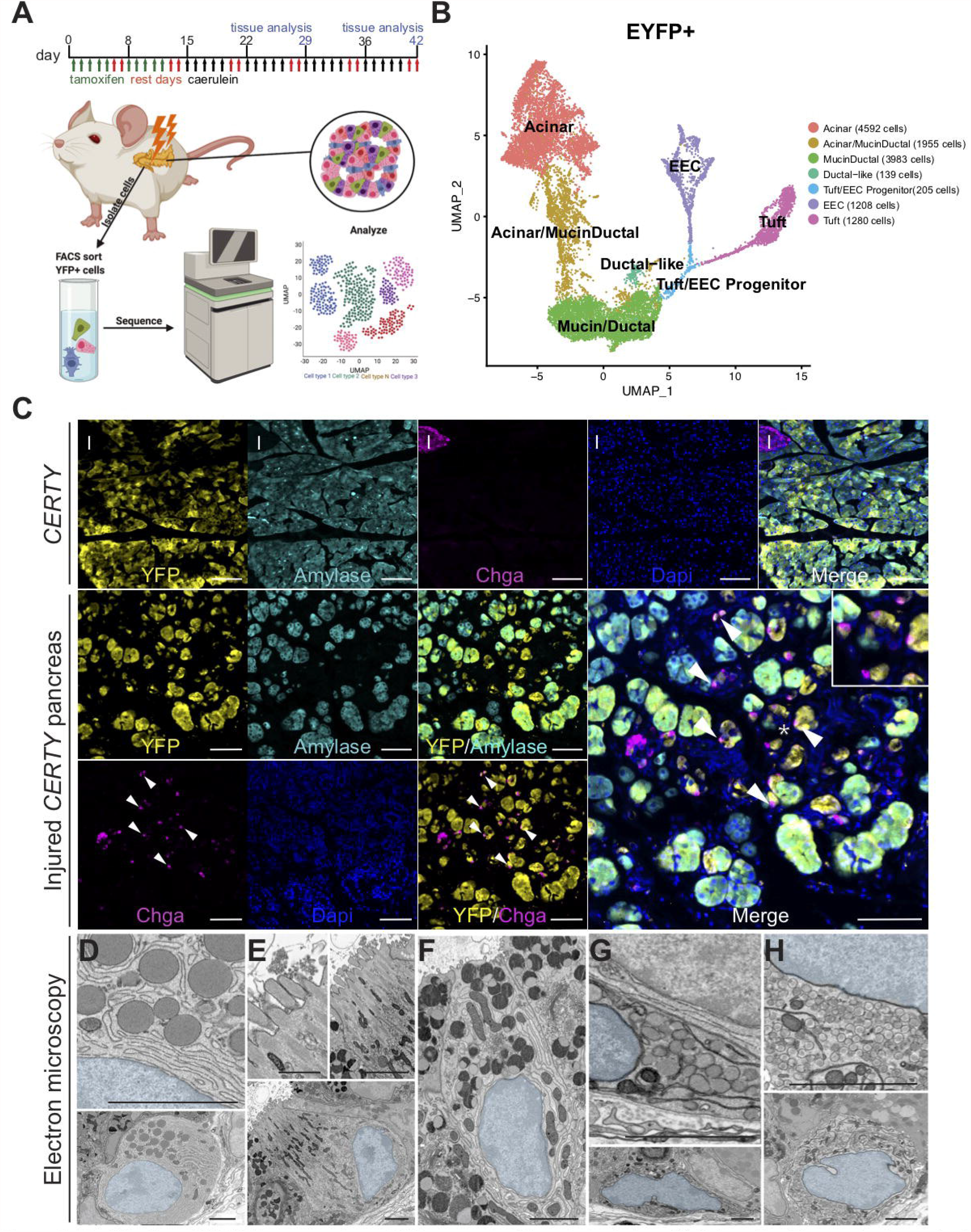
Identification of ADM lineages using scRNA-seq and unbiased clustering. **(A)** Schematic of lineage tracing and scRNA-seq strategy to identify ADM populations. **(B)** UMAP showing annotated clusters in the EYFP+ population. **(C)** Immunostaining for EYFP (yellow), acinar marker amylase (cyan), endocrine marker chromogranin A (CHGA, magenta), and DAPI, blue, in normal and injured *CERTY* pancreata. White arrowheads, YFP+ EECs. Scale bars, 100μm. **(D)** Scanning electron microscopy (SEM) of the injured pancreas highlighting an acinar cell, (E) a tuft cell, **(F)** a mucinous cell, and **(G-H)** two EECs. Scale bars, 500nm; tuft cell inset, 166.7nm.

We removed EYFP-neg stromal cells, potential doublets, and further filtered resulting in 13,362 EYFP+ epithelial cells. Cells from the 4 libraries were integrated using the FastMNN approach (see methods). We identified 16 clusters falling into four broad categories as well as intermediate populations. Acinar cells (*Prss1+Ptf1a+Try4+Cpa1+*; 4592 cells) and tuft cells (*Pou2f3+Siglecf+Trpm5+Gnat3+*; 1280 cells) were identified, consistent with previous studies, validating our methodology^5, 11^. In addition to these populations, we identified mucin/ductal cells (*Muc6+Pgc+Krt19+Car2+*; 3983 cells) and enteroendocrine cells (EECs, *Neurog3+Pax6+Chga+ Neurod1+*; 1208 cells). YFP+ injury-associated acinar cells are dramatically different from normal acinar cells and have upregulated levels of markers related to stress, proliferation, pyloric metaplasia, and ADM (Figure S6A-B, File S4). We also observed a decrease in the percentage of transcripts from digestive enzymes and a several-fold increase in the number of active genes when compared to normal acinar cells (Figure S6C-D)^11, 12^. (Figure S6, File S2). Intermediate ADM-derived populations include acinar/mucin-ductal cells (1955 cells) characterized by decreasing levels of acinar marker genes and increasing expression of mucin/ductal genes, as well as a *Sox4+* EEC/tuft putative progenitor cell population (205 cells) (Figure 1B, S2C-D, File S3).

We used co-immunofluorescence (Co-IF) for EYFP and cell type markers in normal and injured *CERTY* pancreata to validate transcriptomically-identified populations. In uninjured, tamoxifen-treated *CERTY* pancreata, EYFP expression is restricted to acinar cells (amylase+) and is absent from ductal cells and islets (CHGA+), consistent with prior reports^11, 13^. Under conditions of chronic injury, however, EYFP is expressed in both acinar and ductal cells (including tuft cells) as well as a large population of CHGA+ cells consistent with EEC formation (Figure 1C, S3)^5^. As an orthogonal approach to confirm these cells types, we used scanning electron microscopy (SEM) on ultra-thin tissue sections from the pancreata of caerulein-treated mice. SEM readily identified acinar cells by the presence of large zymogen granules and abundant ER. Tuft cells were identified by their distinct microvilli and deep actin rootlets^14^. We identified mucinous cells by mucin granules and several EEC subtypes characterized by electron dense granules (Figure 1D-H). Collectively, these analyses reveal previously unrecognized epithelial heterogeneity in genetically wild type ADM.

### Developmental trajectory analyses identify distinct secretory cell lineages

The cellular heterogeneity identified by scRNA-seq could arise by multiple mechanisms including, but not limited to: 1) direct transdifferentiation from acinar cells, 2) sequential generation from acinar cells to intermediate progenitors, or 3) generation of a common intermediate for tuft and EEC populations, which can then interconvert. We used several computational biology approaches to infer probable lineage trajectories (Figure 2A-D). Trajectory inference algorithms, like Monocle, traverse the collective heterogeneity arising from the asynchronous developmental progression of individual cells^15^. Interestingly, the trajectory inferred by Monocle3 indicates that acinar cells first pass through a mucin/ductal state before forming tuft or EEC lineages (Figure 2A-B). A similar trajectory map is captured through the pCreode algorithm, which describes transitional routes in transcriptional variation in principal component space (Figure 2C, S4A)^16^.

**Figure 2.**
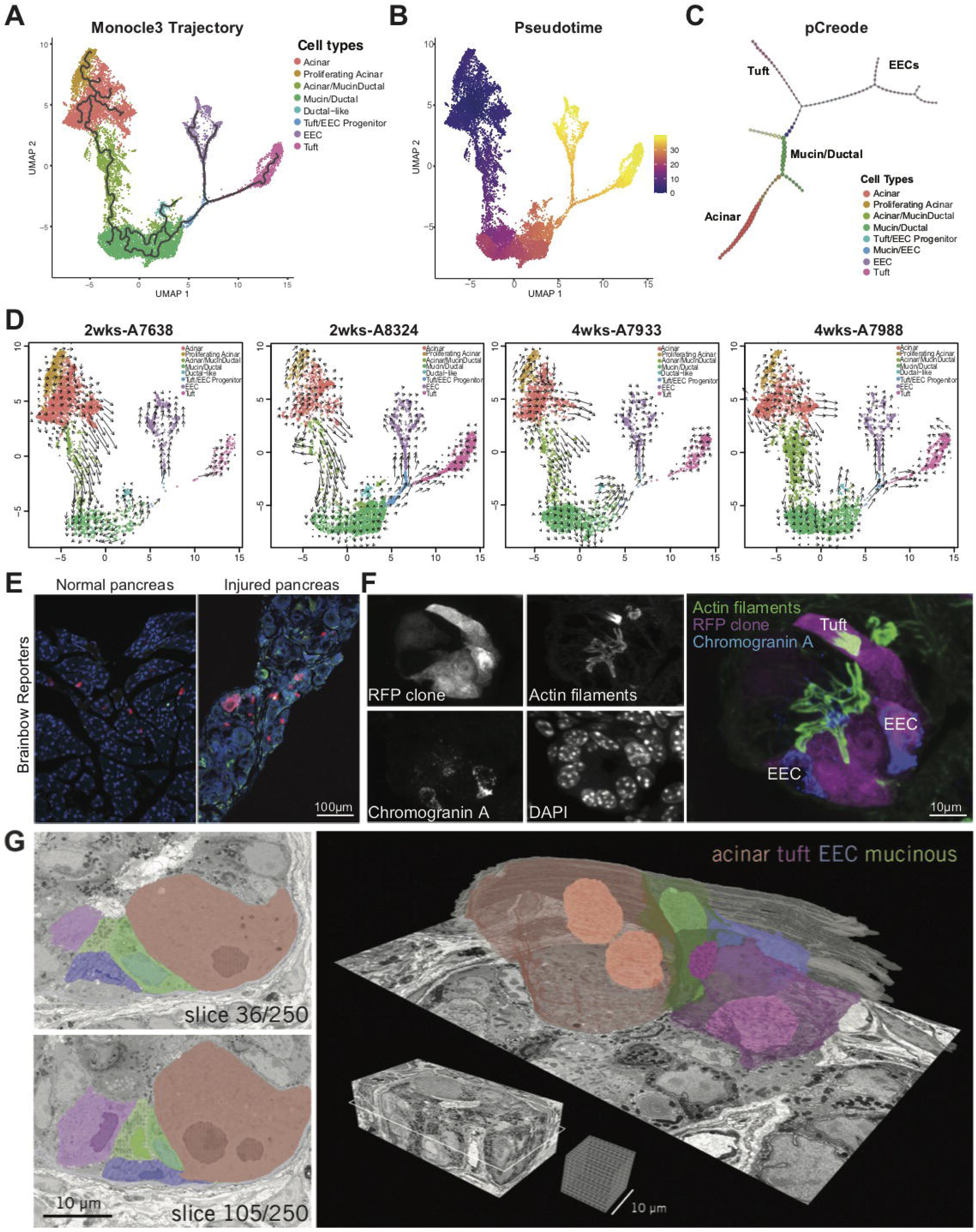
Lineage trajectory analysis of ADM populations. **(A)** Monocle3 trajectory analysis overlaid on the UMAP of EYFP+ cells from injured *CERTY* pancreata. **(B)** Pseudotime projection analysis and **(C)** pCreode approximation of lineage progression of EYFP+ cells. **(D)** RNA velocity analysis of all four datasets. Arrow direction and length indicate the probable lineage trajectory and velocity, respectively. **(E)** Brainbow reporter RFP in untreated and injured pancreata from *Ptf1aCre*^*ERTM/+*^*;Rosa*^*Brainbow*.*2/+*^ mice. Scale bar, 100μm. **(F)** iSIM analysis of a whole RFP+ clone containing a tuft cell (phalloidin) and an EEC (chromogranin A) within the same lesion. Scale bar, 10μm. **(G)** 3-D EM reconstruction of ADM showing a heterogeneous consortium of cell types encapsulated by basement membrane with closely apposed vasculature. From left to right, two partially annotated cross-sections from the 3-D EM volume (volume: 58×29×17.5μm; voxel size: 8-8-70nm, x-y-z). Volumetric reconstructions of cell plasma membranes and nuclei, as well as a portion of the basement membrane rendered in gray, demonstrating the morphological variability between cell types. Scale bars, 10μm.

To independently validate these trajectories, we used RNA velocity, which annotates transitional cell states by leveraging splicing kinetics^17^. We calculated RNA velocity individually in all 4 samples; all samples show a consistent vector field from acinar to mucin/ductal, then branching into EEC and tuft cells (Figure 2D). This trajectory is further supported by the activity of key transcription factors (TFs) (Figure S4B-C). We applied gene regulatory network, or regulon, analysis using the SCENIC package (File S4). Regulon activity scores, generated through AUCell, reflect the enrichment of a TF together with its correlated direct-target genes^18^. We observed a switch from acinar-associated TFs *(Ptf1a+Rbpjl+Bhlha15/Mist1+)* to Mucin *(Spdef+Creb3l1+Creb3l4+)* and Ductal *(Sox9+Hnf1b+Onecut1+)* associated TF activation in the Acinar/MucinDuctal intermediate population (Figure S4B-C)^19-21^. Collectively, these data are consistent with a model in which injured acinar cells form a mucin/ductal progenitor population that seeds tuft cells and EECs as distinct lineages.

We tested this hypothesis by conducting lineage tracing with a Brainbow reporter. Cre recombination in *Ptf1a*^*CreERTM*^*;Rosa*^*Brainbow*.*2/+*^ mice results in the random expression of 1 of 4 reporters (YFP, GFP, RFP, CFP) in acinar cells. Mice were first treated with a very short course of tamoxifen followed by caerulein (as shown in Figure 1A). Immunofluorescence (IF) analysis of uninjured pancreata shows solitary cells expressing Brainbow reporters. Injured pancreata, in contrast, are characterized by reporter expression in partial or entire lesions, demonstrating clonal expansion from single acinar cells (Figure 2E, S4D-E). To examine cellular heterogeneity, we conducted co-IF and on whole clones stained for markers of tuft cells (phalloidin, labels the microvilli and actin rootlets) and EECs (CHGA) and identified both cell types residing within the same clones (Figure 2F). To further evaluate cellular heterogeneity within individual ADM lesions, we performed SEM on 250 serial sections taken of a single metaplastic duct. Segmentation and 3-dimensional reconstruction of the processed image stack revealed an acinar cell adjacent to mucinous cells, a tuft cell, and an EEC with delta cell morphology (Figure 2G)^22^. Interestingly, the identified acinar cell is binuclear; how this factors into acinar cell plasticity is currently unknown, but it has been shown that binucleate cells are resistant to cell cycle entry^23^.

### Injury-induced ADM as a pyloric-type metaplasia

Top differentially expressed genes (DEGs) in the mucin/ductal population include *Muc6, Tff2*, and *Gif* which are characteristic of SPEM in the stomach (Figure 3A, S2D,File S3)^24^. SPEM is a process similar to ADM where digestive-enzyme-secreting chief cells undergo metaplasia in response to injury or oncogenic mutation^25, 26^. SPEM occurs within the overall reorganization --known as pyloric metaplasia --characterized by gastric units in the body (main portion) of the stomach reprogramming to resemble the distal (pyloric) region^25, 27, 28^. We identified a number of canonical SPEM markers *(Cd44, Aqp5, Gkn3, Dmbt1, Wfdc2)* enriched in our mucin/ductal population as well as TF Gata5, a transcription factor that is widely expressed in gastric/intestinal epithelium but not in normal pancreas epithelium (Figure 3A, S5A). We also identified markers of chief cells *(Gif* or *Cblif, Pga5, Pgc)*(Figure 3A, S5B). To confirm these findings, we conducted immunohistochemistry (IHC) and/or Co-IF for CD44v9, GKN3, AQP5, WFDC2, and GIF. Consistent with our sequencing data, we identified the expression of SPEM and chief cell markers in injured pancreata, whereas expression is absent from the normal pancreas (Figure 3B, S5C). To confirm that this phenotype is ADM-specific, we compared our mucin/ductal population to normal pancreatic ductal scRNA-seq datasets and found SPEM signatures to be restricted to ADM (Figure S6E-F, File S2)^11, 29^. Finally, we conducted SEM on ADM and identified cells with admixed mucin and digestive enzyme-containing granules, similar to the phenotype of SPEM associated with gastric injury (Figure 3C).

**Figure 3.**
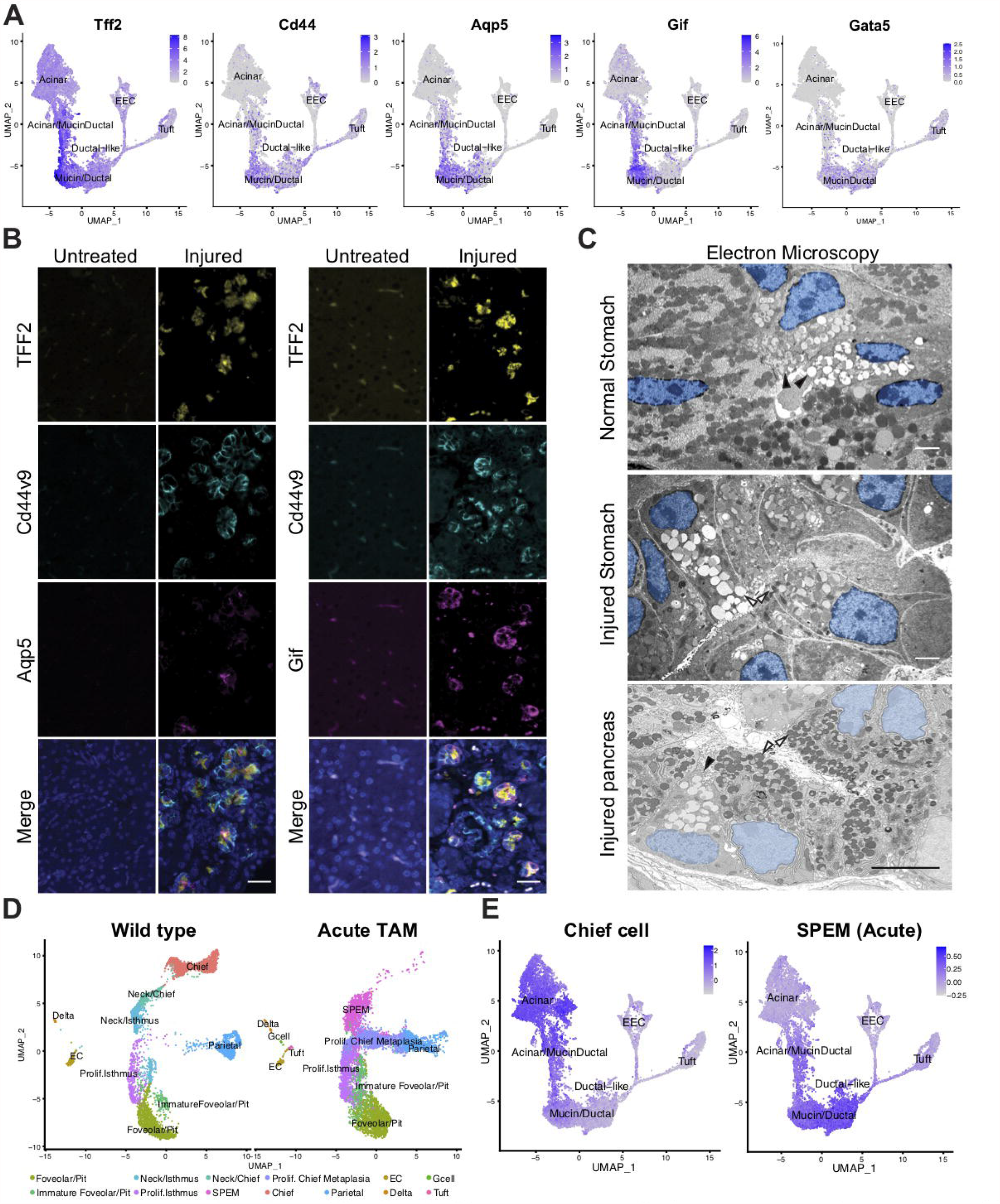
ADM as a pyloric-type metaplasia. **(A)** Expression of SPEM marker genes *Tff2, Cd44*, and *Aqp5*, chief cell marker gene *Gif*, and TF *Gata5* overlaid on the UMAP of EYFP+ cells. **(B)** Co-IF for SPEM markers TFF2 (yellow), CD44v9 (cyan), and AQP5 or GIF (magenta) and dapi (blue). Scale bars, 25μm. **(C)** EM of mature mucus neck cells (black arrowheads) in the normal stomach, SPEM in an acute model of gastric injury (white arrowheads), and mucin/ductal cells from pancreatic ADM. Analogous granules in ADM are marked by arrowheads. Scale bars, 2μm. **(D)** UMAP of scRNA-seq data collected from normal murine stomach and a SPEM model of acute gastric injury, from Bockerstett et al. **(E)** Expression of chief cell and acute SPEM signatures overlaid on the UMAP of EYFP+ cells. Gene signatures generated from **(D)**. EC, enterochromaffin; Prolif, proliferating.

Based on these findings, and prior studies noting several of these markers in PanIN, we hypothesized that acinar cells undergo a metaplasia within the general rubric of pyloric metaplasia with specific parallels to SPEM^11, 30^. To test this hypothesis, we compared our ADM dataset to a recently published scRNA-seq study by Bockerstett et al., comparing normal gastric epithelium to that in several models of SPEM (Figure 3D and S7A-C)^31^. When we overlaid DEGs from normal gastric cell types onto our EYFP+ dataset, we found the chief cell transcriptome to be most closely related to pancreatic acinar cells (Figure 3E, S7A, File S5). To generate SPEM signatures, we identified DEGs between SPEM populations and normal chief cells in either a mouse model of acute gastric injury (73 genes) or of chronic autoimmune injury (54 genes) and overlaid these signatures on our ADM dataset (File S6). We found both SPEM signatures to be enriched in our mucin/ductal population (Figure 3E, S7C).

We also identified TFs common to ADM and SPEM. Gata5 and several mucin related TFs *(Spdef, Creb3l1* and *Creb3l4)* are enriched in the mucin/ductal cluster (Figure S4B, S7D)^19, 32, 33^. To determine if these transcriptional programs are also activated in SPEM, we examined regulon module activity in both the acute and chronic SPEM scRNA-seq datasets^18^. We found all the identified regulons elevated in SPEM suggesting similar regulatory programs in both disease states (Figure S7E-F).

### ADM results in EEC heterogeneity

We next explored heterogeneity within our lineage traced Tuft/EEC population. Re-clustering of the Tuft+EEC subset revealed 9 cell clusters separated into Tuft/EEC progenitors, EECs, or tuft lineages (Figure 4A, File S7). Trajectory inference using Monocle3 and RNA velocity suggests that EEC and tuft cell lineages originate from a *Sox4+* common progenitor population (Figure 4B-C, 4F), which has previously been reported in the intestines^34^. Within the EEC population, we identified 5 distinct clusters including a progenitor-like cluster (*Neurog3+*, Figure 4D), and putative enterochromaffin *(Ddc+Tac1+Chgb+)*, delta *(Sst+)*, epsilon *(Ghrl+)*, and PP/gamma cells *(Ppy+)* (Figure 4E, S8A)^13^. Gast (gastrin) expression is detected in a subset of *Ddc+Tac1+* cells; Cck (cholecystokinin) in a subset of *Ghrl+* cells. Insulin is not detected; minor glucagon expression is present within the *Ppy+* cluster (Figure S8A-B)

**Figure 4.**
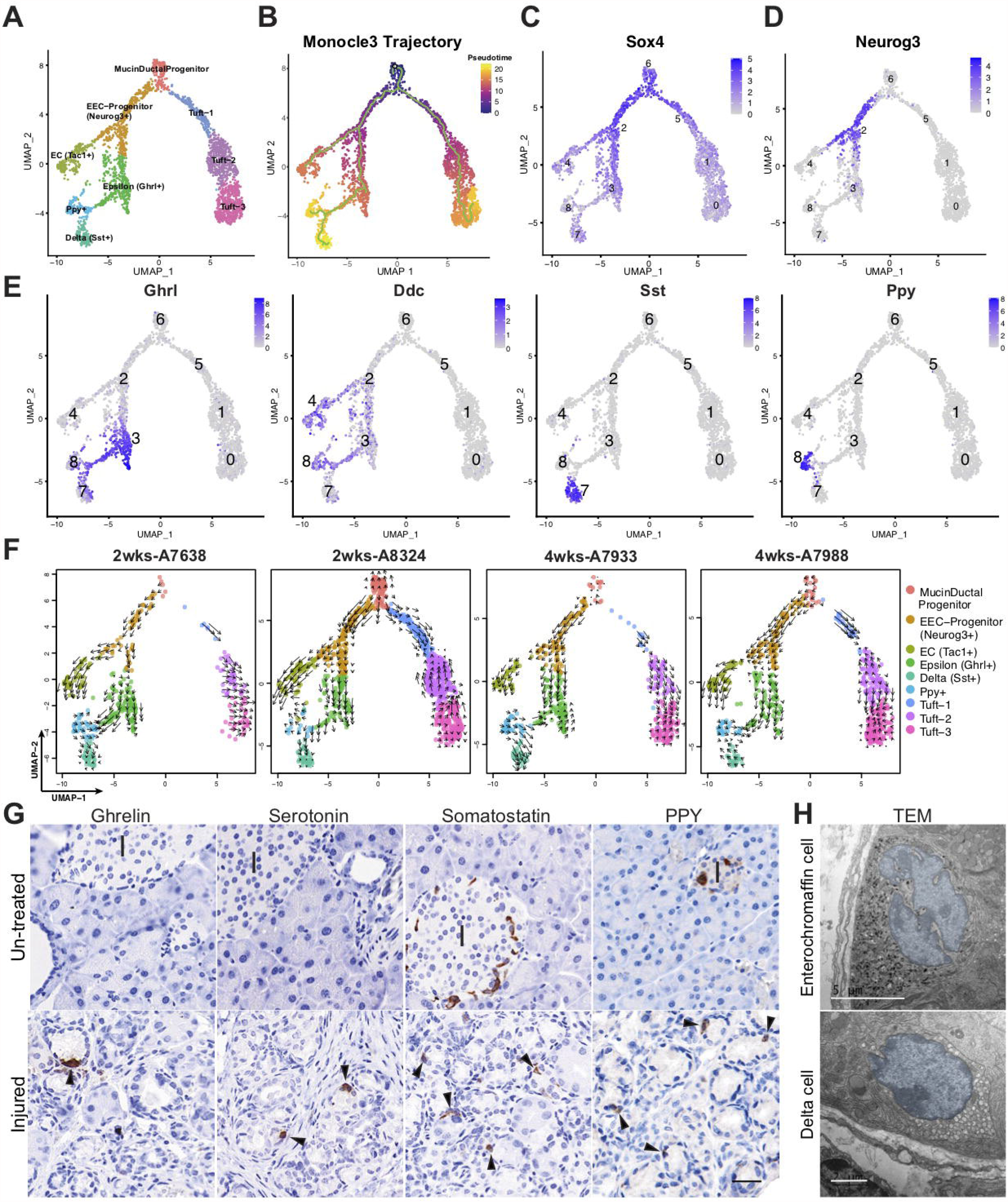
ADM results in substantial enteroendocrine cell heterogeneity. **(A)** UMAP of cell clusters identified in the combined tuft and EEC dataset identified in Figure 1B and labeled by cell type. **(B)** Monocle3 trajectory analysis overlaid on the UMAP in (A). **(C)** Expression of TFs *Sox4* and **(D)** *Neurog3* overlaid on the UMAP from (A). (E) Expression of epsilon cell marker *Ghrl*, enterochromaffin cell marker *Ddc*, delta cell marker, *Sst*, and PP/gamma cell marker *Ppy* overlaid on the UMAP from (A). (F) RNA velocity analysis of all four datasets. Arrow direction and length indicate probable lineage trajectory and velocity, respectively. (G) IHC for ghrelin *(Ghrl)*, serotonin (generated by *Ddc*), somatostatin *(Sst)*, and pancreatic polypeptide *(Ppy)* in untreated and injured pancreata. Black arrows, positive cells. I, islets. Scale bar, 25μm. (H) TEM of an enterochromaffin cell or a delta cell in the injured pancreas. Scale bars, 5 and 2μm respectively.

To confirm expression of these hormones at the protein level, we performed IHC and detected ghrelin, serotonin (*Ddc* and *Tph1*), somatostatin, pancreatic polypeptide, and gastrin in injury-induced ADM (Figure 4G, S8F). To determine if these hormone-producing EECs are truly associated with ADM and not islets, we conducted lineage tracing with *CERTY* mice and identified co-expression of EYFP in all 4 EEC subtypes (Figure S8G). Analysis of EM conducted on injured pancreata identified cells with enterochromaffin cell (serotonin-producing, pleomorphic granules) and delta cell (somatostatin-producing, round granules) morphology (Figure 4H)^35^.

To identify putative regulatory processes involved in EEC subtype specification, we examined the activity of key TFs and signaling molecules. We found that the *Neurog3+* ECC progenitor state is associated with expression of Notch pathway inhibitors and subsequent branching into enterochromaffin and epsilon cell lineages is associated with changes in key TFs^36^. Enterochromaffin lineage formation correlates with *Nkx6-1* activation and a switch from *Pax4* to *Pax6* expression^37^. The epsilon lineage is associated with activation of *Arx* and *Isl1*. Subsequent branching into delta and PP/gamma cells is accompanied by reactivation of Pax6. Delta cells are characterized by a lack of *Arx*, reactivation of *Sox9*, and expression of *Pou3f1* (Figure S8C-E) ^38,39, 40^.

### Transcriptomic changes associated with tuft cell development

A scRNA-seq study of intestinal epithelium previously demonstrated that tuft cells can be categorized as either neuronal or inflammatory-like based on marker expression^41^. To identify tuft sub-populations in our dataset, we conducted DEG and GO Term analysis of tuft cell clusters (6, 5, 1, 0) including a total of 1280 cells (Figure 5A, S9A). Consistent with RNA velocity results (Figure 4F), we observed a transition from a *Sox4+* progenitor population (cluster 6) to a *Pou2f3+Spib+* population (cluster 5 and cluster1); *Pou2f3* is the master regulator for tuft cell formation (Figure 5A-C)^42^. Subsequent clusters 1 and 0 are enriched for known tuft cell markers and signaling molecules (i.e. *Trpm5, Il25*)(Figure 5A, 5J), suggesting that our dataset reflects the transcriptomic changes associated with tuft cell development, rather than separate, functional subtypes.

**Figure 5.**
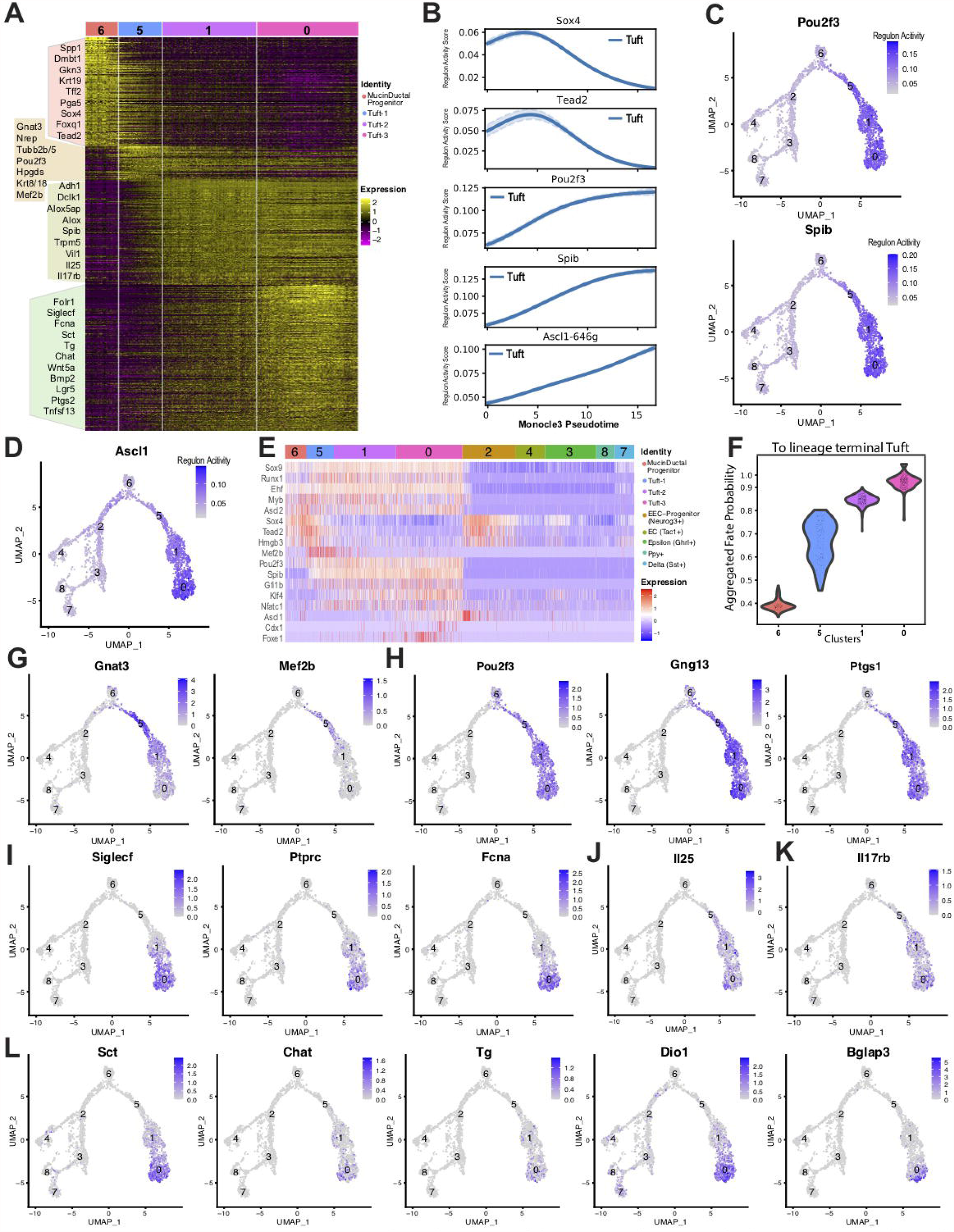
Transcriptomic changes associated with tuft cell formation. **(A)** Heatmap of scaled DEGs for the 4 clusters of the tuft cell lineage. A selection of known marker genes are labeled on the left. Cells were ordered by increasing pseudotime within each cluster group. **(B)** Smoothed regulon activity score trend of the tuft cell lineage in Monocle3 pseudotime. **(C-D)** Regulon activity scores of *Pou2f3, Spib*, and Ascl1 overlaid on the Tuft+EEC UMAP. **(E)** Heatmap of scaled RNA expression of tuft-related TFs. Cells are ordered by increasing pseudotime within each cluster group. **(F)** The aggregated fate probability towards terminal tuft lineage formation. **(G-K)** Expression of select genes overlaid on the Tuft+EEC UMAP, labeled by Seurat cluster.

To test this hypothesis, we conducted regulon analysis and identified candidate TFs and down-stream regulatory networks associated with tuft cell development^18^. We examined regulon activities as a function of Monocle3 Pseudotime, moving from cluster 6 -> 5 -> 1 -> 0. Consistent with RNA velocity and DEG analysis, we identified a decrease in Sox4 activity, and an increase in Pou2f3, Spib, and Ascl1 activity (Figure 5B-D). Additional regulators include neuronal development-associated TFs *(Tead2, Hmgb3, Mef2b)*, TFs associated with hematopoietic differentiation and inflammatory cell function *(Runx1, Myb, Gfi1b, Nfatc1*, and *Foxe1)* and TFs associated with epithelial differentiation *(Ehf, Klf4)* (Figure 5E)^43-45^. We next examined cell fate probability using a directed single-cell fate mapping approach. Cells were decomposed into macrostates matching our seurat clusters (Figure S9B). Directed fate probabilities reveal a continuous process of tuft cell formation, which is shown by a steady gain in tuft cell lineage fate probabilities moving from cluster 6 -> 0, consistent with tuft cell development and maturation (Figure 5F, S9C). This is consistent with transcriptional activity changes in many tuft cell marker genes. The first group we identified includes genes enriched in cluster 5, where expression fades throughout development *(Gnat3, Mef2b, Nrep)*(Figure 5G, S9E). The second group represents a steady level of gene expression throughout the pseudotemporal ordering from cluster 5 to cluster 0 *(Pou2f3, Gng13, Ptgs1, etc*.)(Figure 5H, S9F). The last group includes genes associated with the late stage of tuft cell formation where expression increases along tuft cell development *(Siglecf, Ptprc, Fcna)*(Figure 5I, S9G).

Finally, we explored the tuft cell classifications previously reported in small intestines in our dataset by overlaying consensus gene signatures^41^. We found that the inflammatory signature is enriched late in tuft cell development, while the neuronal signature is enriched at a relatively early stage and remains expressed (Figure S9D). Recently, Manco et al., used ‘clump sequencing’ to identify spatial expression programs of intestinal secretory cells^46^. The authors found that tuft cells associated with intestinal crypts, where cells are more stem-like, express *Sox4* and are enriched for the neuronal tuft gene signature. Tuft cells associated with the intestinal villus, where cells are most differentiated, are enriched for functional genes like *Il17rb, Fapb1* and the inflammatory tuft signature. Consistent with this, we identified expression of *Il17rb, Fabp1*, and a number of other secretory signaling molecules and hormones enriched in differentiated tuft cell cluster 0 (Figure 5J-L, S9G). Collectively, these data identify regulatory programs and gene expression changes associated with tuft cell maturation.

### *Kras*^*G12D*^ expression drives PanIN-specific changes in ADM

ADM occurs in response to oncogenic *Kras*^*G12D*^ and may serve as a precursor lesion for PDAC. Recently, Schlesinger et al., published a scRNA-seq study of pancreatic tumorigenesis including lineage tracing using *LSL-Kras*^*G12D*^*;Ptf1a*^*CreERTM*^*;LSL-tdTomato (KCT)* mice^11^. To identify similarities and differences between injury and oncogene-induced ADM, we compared EYFP+ cells (13,362 cells) to *tdTomato+* cells FACS-collected from a 6-month-old *KCT* mouse (2719 cells, Figure S10A). We integrated the two datasets using fast MNNand identified 18 distinct clusters (Figure S10B-C). Using the aforementioned gene signatures as well as those in Schlesinger et al., we found a number of populations common to both disease states (e.g. tuft and EECs, Figure 6A). Metaplasia transcription factors *Foxq1* and *Onecut2* (described in Schlesinger et al.) are also expressed in injury-induced ADM (Figure S10D-E)^11^. Acinar populations in *Kras*^*G12D*^ ADM are enriched with regenerating (REG) protein family members (*Reg2+Reg3a+Reg3g+* ‘Acinar-REG+’), while those associated with injury-induced ADM express higher levels of ribosome transcripts; consistent with the recently identified ‘secretory acinar cell’ phenotype (‘Acinar-S’) thought to function primarily in production of digestive enzymes (Figure S10F)^8^.

**Figure 6.**
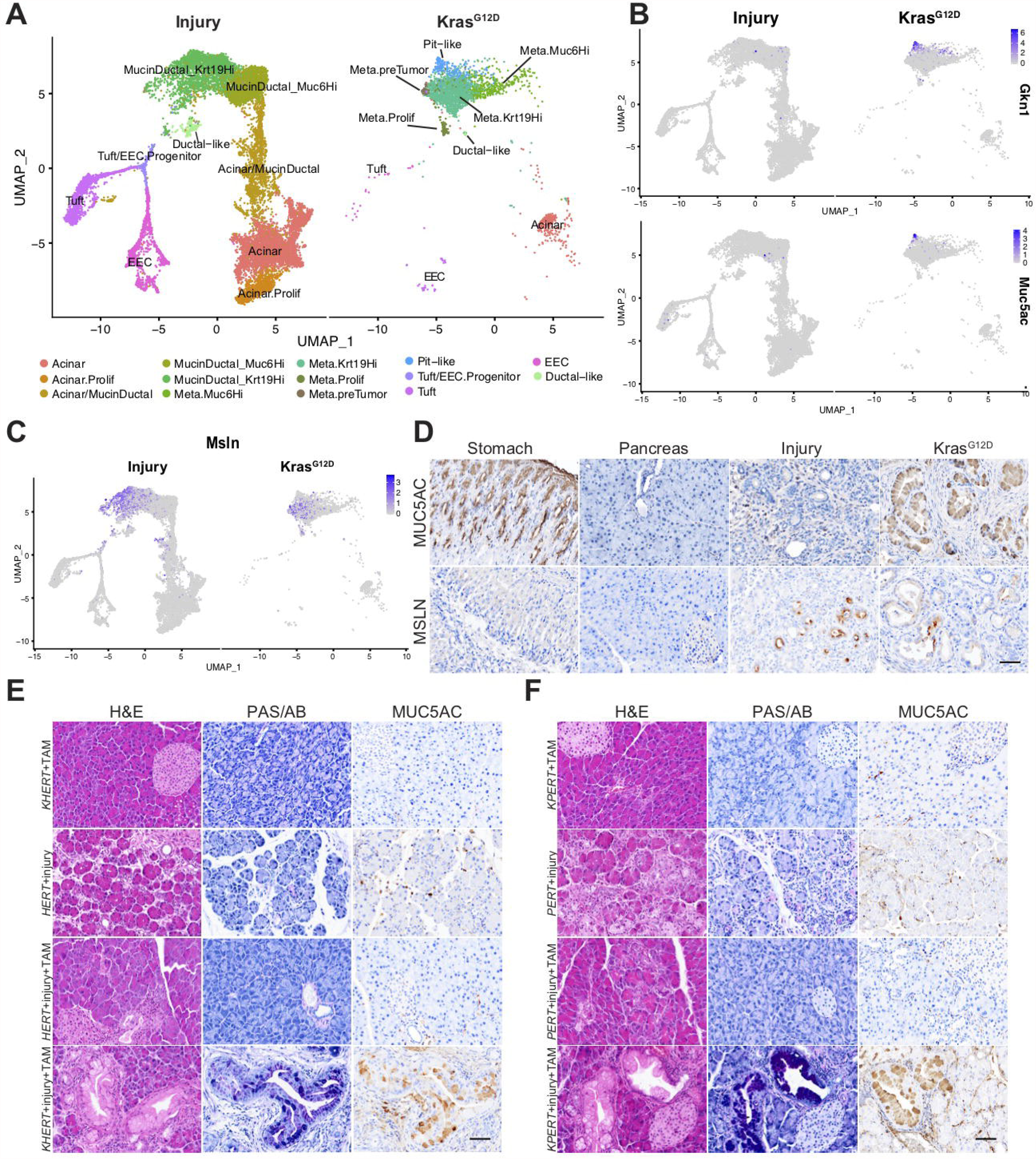
*KrasG12D* expression drives PanIN-specific changes in ADM. **(A)** UMAP of FastMNN integrated datasets from the injury-induced ADM scRNA-seq dataset and an oncogene-induced ADM dataset from a six-month-old *KCT* mouse derived from Schlesigner et al. **(B)** Expression of pit cell markers *Gkn1* and *Muc5ac* and **(C)** pre-tumor marker *Msln*. **(D)** IHC for MUC5AC or MSLN in the normal stomach or pancreas as well as in injury or oncogene *(Kras*^*G12D*^)-induced ADM and neoplasia. **(E-F)** H&E, PAS/AB (mucins) or MUC5AC expression in *KHERT* or *KPERT* mice treated with tamoxifen, *HERT* or *PERT* mice treated with caerulein or caerulein followed by tamoxifen, and *KHERT* and *KPERT* mice first treated with caerulein followed by tamoxifen. Scale bars, 50μm.

Differences between the ADM populations in the two datasets include a lack of senescence markers, and ‘metaplastic pre-tumor’ populations in the injury model (Figure 6A)^11^. Further, we identified distinct differences in the mucin/ductal population between the two disease states including a shift from *Muc6*-high,*Tff2*-high,*Krt19*-low cells to *Muc6*-low,*Tff2*-low,*Krt19*-high cells in the KCT pancreas, possibly reflecting enhanced disease progression (Figure 6A and S10B). DEG analysis of the mucin/ductal cell clusters between the two disease states identified significantly higher expression of SPEM markers in injury-induced ADM, consistent with the presence of a larger *Muc6*-high,*Tff2*-high,*Krt19*-low population. Oncogene-induced ADM, in contrast, is characterized by significantly higher expression of proto-oncogenes such as *Areg, Fos* and *Junb*, known down-stream targets of *Kras*^*G12D*^ (Figure S10G)^47^. Schlesinger et al., described the formation of a gastric pit cell-like population *(Tff1+Gkn1+Muc5ac+)* in oncogene-induced ADM (Figure 6A-B)^11^. To assay injury-induced ADM for pit cells, we conducted IHC for MUC5AC and found expression to be specific to *Kras*^*G12D*^ pancreata. Interestingly, a number of pre-tumor markers described by Schlesinger et al., *(Msln, Mmp7, Marckls1*, and *Igfbp7)* are, in fact, expressed in injury-induced ADM (Figure 6C-D, S10H-I)^48^. To confirm expression of these pre-tumor markers, we conducted IHC for MSLN and detected positive cells within the epithelium of injured pancreata (Figure 6D). These data suggest that ‘pre-tumor’ cells represent a transitional state co-opted by oncogenic *Kras* during tumorigenesis.

To determine if *Kras*^*G12D*^ expression is sufficient to drive the aforementioned differences in injury-versus oncogene-induced ADM, we used several inducible Cre drivers to target *Kras*^*G12D*^ expression to ADM populations. *Hnf1b:CreER*^*T2*^ mice have been described previously^49^. In the adult pancreas, Hnf1b expression is restricted to ductal cells and Bailey et al., have shown that two alleles of *Trp53*^*R172H*^ in addition to *Kras*^*G12D*^ are required to induce transformation^50^. *Kras*^*G12D*^ expression alone results in no discernable phenotype in the pancreas. To determine if *Kras*^*G12D*^ expression in injury-induced *Hnf1b+* ductal cells is sufficient to drive PanIN-associated changes, we bred *Kras*^*G12D*^*;Hnf1b:CreER*^*T2*^*;Rosa*^*LSL-YFP/+*^ *(KHERTY)* mice, induced injury and ADM with caerulein first, and then activated *Kras*^*G12D*^ expression after ADM had formed (Figure S11A-C). We found that, while the *Hnf1b:CreER*^*T2*^ pancreas heals after injury and tamoxifen treatment, that targeting *Kras*^*G12D*^ expression to ADM results in 100% (n=6) of *KHERTY* pancreata forming MUC5AC+ PanIN (Figure 6E).

As the *Hnf1b:CreER*^*T2*^ mouse model drives *Kras*^*G12D*^ expression from both normal and ADM-induced ductal cells, we next conducted this experiment using ADM-specific *Pou2f3-cre*^*Ert2*^*;Rosa*^*mT/mG*^ mice^51^. We have previously shown that POU2F3 expression is absent from the normal pancreas, that 99% of POU2F3+ tuft cells are derived from ADM, and that tuft cells are rare in the injured pancreas (making up ∼0.6% of the epithelium)^5, 7^. As shown in Figure 5H, Pou2f3 is expressed early in injury-induced EEC/tuft specification. To determine if ADM-induced Pou2f3+ cells are susceptible to oncogenic transformation and if *Kras*^*G12D*^ expression is sufficient to drive the changes identified in our scRNA-seq analyses, we bred *Kras*^*G12D*^; *Pou2f3-cre*^*Ert2*^*;Rosa*^*mT/mG*^ (KPERT) mice. We found that the *Pou2f3-cre*^*Ert2*^ pancreas heals after recovery from injury and tamoxifen treatment, however, 40% (4/10 mice) of KPERT pancreata contained MUC5AC+ PanIN (Figure 6F, S11D-F). PanIN are sparse in this model, consistent with the rarity of POU2F3 positive cells in injury-induced ADM. Collectively, these data identify changes in the epithelium between injury and oncogene-induced ADM and demonstrate that expression of *Kras*^*G12D*^ in injury-induced ADM is sufficient to drive these changes.

### Human pancreatic metaplasia as a pyloric-type transition

To determine if the epithelial composition of human chronic pancreatitis ^52^ reflects the diversity identified in mouse models, we evaluated marker expression in single nucleus RNA sequencing (sNuc-seq) data collected from human pancreata and conducted immunofluorescence on tissue samples. Recently, Tosti et al., generated a comprehensive human pancreas cell atlas consisting of ∼113,000 nuclei from healthy adult donors and 2,726 nuclei from two patients with chronic pancreatitis^8^. Consistent with mouse data, the pancreatitis dataset reflects an increase in a mucin/ductal population (*MUC5B+/KRT19+;* “MUC5B+Ductal”) and the appearance of tuft cells, as compared to the normal pancreas (Figure 7A, S12A). To directly compare the human and murine mucin/ductal populations, we overlaid their respective gene signatures on their counterpart species dataset and found enrichment in the Mucin/Ductal clusters (Figure S12B).

**Figure 7.**
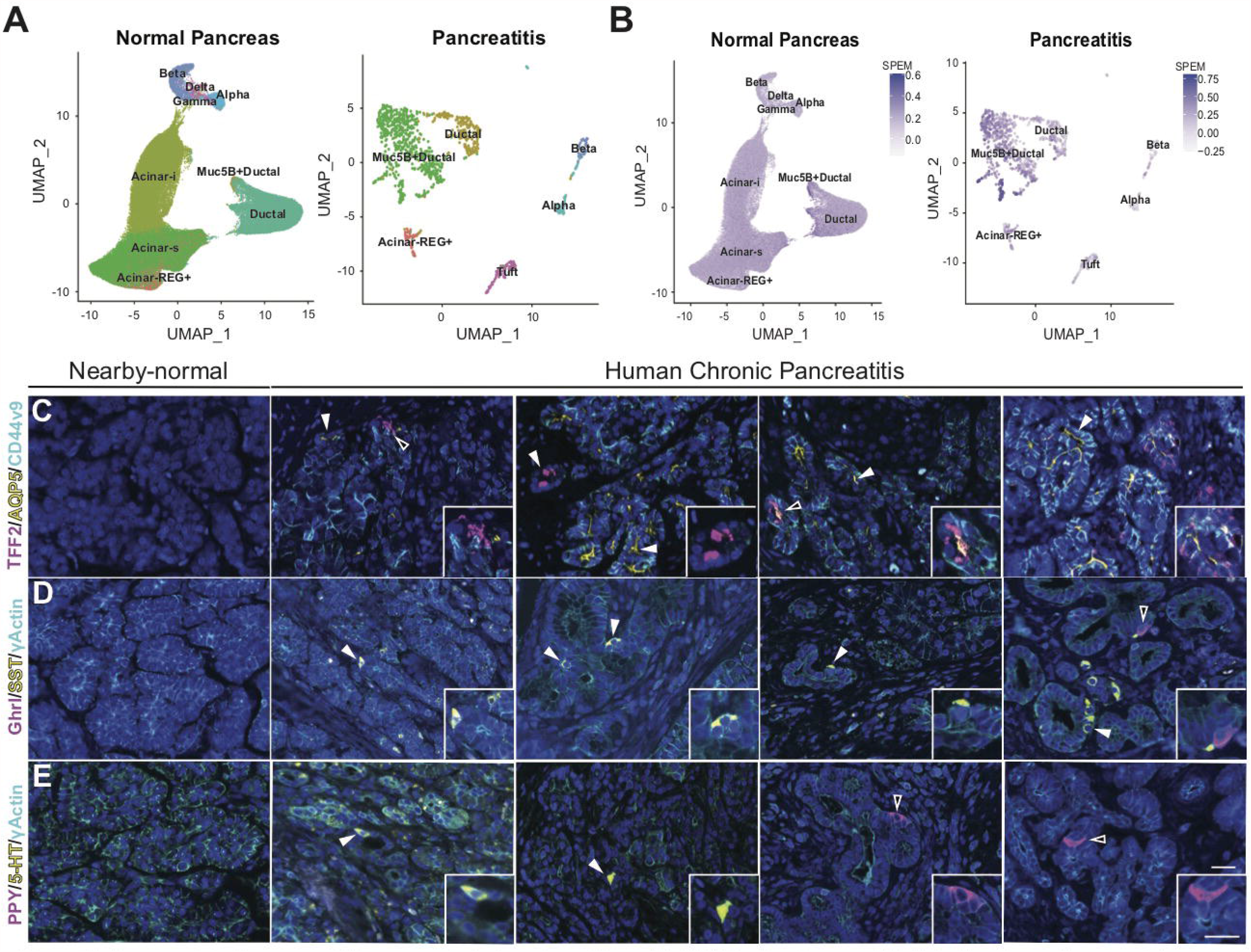
Human chronic pancreatitis as a pyloric-type transition. **(A)** UMAPs showing annotated clusters from sNuc-seq data collected from normal human pancreata (∼113k nuclei) or the injured pancreata of two patients with chronic pancreatitis (2,726 nuclei) derived from Tosti et al. **(B)**. Expression of the humanized acute SPEM signature (Figure 3D) overlaid on the UMAPs from (A). (C) Co-IF for SPEM markers (TFF2, magenta; AQP5, yellow; CD44v9, cyan), **(D)** epsilon and delta cell markers (Ghrl, magenta; SST, yellow; γActin, cyan), and **(E)** gamma and enterochromaffin cell markers (PPY, magenta; 5-HT, yellow; γActin, cyan). Dapi, blue. Representative images from 4 different patients are shown. Scale bars, 25μm.

To determine if human ADM represents a pyloric-type metaplasia transition, we humanized and overlaid the murine acute and chronic SPEM signatures on the human pancreatitis dataset. In both analyses, we found enrichment of the SPEM signature in the MUC5B+Ductal population (Figure 7B, S12C). Analysis of individual markers identifies *TFF2, WFDC2*, and *CD44* (Figure S12A). To validate these findings, we conducted immunostaining on pancreata from 15 patients with chronic pancreatitis, pre-screened by a pathologist for the presence of metaplasia, but varying in terms of etiology and severity. First, we evaluated the expression of SPEM markers TFF2, AQP5, and CD44v9 (generated by *CD44*). As compared to normal pancreas, we identified up-regulation of CD44v9 and AQP5 in acinar cells in areas of tissue damage, in all 15 samples (Figure S12D). Occasionally, TFF2 was co-expressed in these regions (triple positive clusters present in 13 of 15 samples), but was more often found in metaplastic structures lacking acinar morphology, in PanIN-like lesions, and in duct glands associated with large ducts (Figure 7C).

We next evaluated the presence of tuft cells and EECs in human chronic pancreatitis. Consistent with prior reports, we identified tuft cells by expression of marker phospho-Girdin in areas of metaplasia associated with injured acini, in normal ducts, and in incidental PanIN (10/15 patient samples, Figure S12E)^6, 53^. When we interrogated the sNuc-seq dataset for tuft cell TFs identified in our EYFP+ dataset, we found expression of *SOX4, POU2F3, SPIB*, and *RUNX1* in human tuft cells (Figure S12F). To assess EEC heterogeneity, we conducted IF for ghrelin (Ghrl), somatostatin (SST), pancreatic polypeptide (PPY), and serotonin (5-HT). All four hormones could be detected in islets, though the proportions of each varied by patient. All hormones were associated with aberrant ductal structures, rather than injured acini, consistent with formation later in ADM. Ghrelin and PPY expression is rare in metaplasia, but both can be found in large ducts. 5-HT and SST are expressed in metaplasia, PanIN-like lesions, and in intralobular ducts (Figure 7D-E). Finally, we assessed the expression of pre-tumor markers in the pancreatitis sNuc-seq dataset. We found expression of *MMP7* and *MARCKSL1*, but not *IGFBP7*, in MUC5B+Ductal cells, though it is not specific. *MSLN*, though, could be detected only in MUC5B+Ductal cells (Figure S12G).

Overall, the staining patterns in humans are largely reminiscent of the murine model, including the formation of distinct cell types (mucin/ductal, tuft, EECs) and expression of SPEM signatures. Differences include the lack of TFF2 in injured acini and expression of individual genes (i.e. *Muc6* vs. *MUC5B*). Further studies are required to determine if these differences reflect the heterogeneity of etiology, chronicity and severity of CP in patients selected to undergo surgical resection.

## DISCUSSION

Using lineage tracing, scRNA-seq, multiple computational biology approaches (that rely on non-overlapping assumptions), immunostaining, and high-resolution ultramicroscopy, we have identified significantly more epithelial heterogeneity in injury-induced pancreatic ADM than previously appreciated. Brainbow lineage tracing demonstrates that acinar cells can clonally expand, forming secretory types identified in these analyses. While scRNA-seq provides information regarding cell type markers, high-resolution ultramicroscopy is critical to the identification of these populations as it provides structural information, corroborating the formation of defined cell types. Collectively, these data are consistent with a model by which acinar cells undergo a pyloric type metaplasia resulting in mucinous ductal cells; it is from this state that differentiated tuft cells or EECs emerge.

We have discovered that chronic injury-induced ADM results in substantial EEC heterogeneity. ADM-derived EEC formation was previously described by Farrell et al., in PanIN resulting from expression of oncogenic *Kras*^*G12D*^ by lineage tracing and expression of marker synaptophysin^54^. The term ‘enteroendocrine cell’, however represents a lineage populated by multiple subtypes characterized by the expression, or co-expression, of various hormones. While best known for regulating intestinal function, insulin secretion, nutrient assimilation and food intake in normal tissues, evidence points to poorly understood roles for several of these hormones in inflammation and tumorigenesis^35, 55^. In particular, somatostatin is known to have diabetogenic properties, which, when combined with the observation that adult-onset diabetes is a known association of pancreatic adenocarcinoma, suggests the intriguing idea that tumor-derived EEC have an important role in the endocrine/metabolic symptoms associated with this malignancy^56^. These data underscore the importance of understanding the contribution of EECs, including their regulatory processes and secretory products, in the contexts of pancreatic injury and tumorigenesis.

Altogether, we propose that this carefully orchestrated process of acinar cell de-and trans-differentiation exists to mitigate injury and may invoke a subset of a generalized pyloric metaplasia response that can occur in many organs in the GI tract^28^. Ever more evidence is accumulating that there are shared patterns of regenerative changes that depend on cellular plasticity. Here, we observe striking transcriptional overlap between metaplastic cells in the pancreas and those of the stomach. Our results are in line with recent studies indicating that mature cells like gastric chief cells and pancreatic acinar cells have a shared injury-induced plasticity program, paligenosis, that governs their conversion from the differentiated to regenerative states^3, 25, 57^. Cycles of paligenosis that fuel pyloric metaplasia may allow cells to accumulate mutations and increase the risk for progression to cancer.

Interestingly, many of the epithelial populations identified in injury-induced ADM, such as tuft and EECs, are shared by oncogenic *Kras*^*G12D*^-induced ADM, suggesting that they represent an early event in tumorigenesis. It is the differences between the two disease states, however, that may provide important clues as to which population(s) signal transformation or progression to cancer. For example, the formation of pit-like cells *(Muc5ac+Gkn1+Tff1+)* is specific to *Kras*^*G12D*^-induced ADM^11^. Here, we demonstrate that activation of *Kras*^*G12D*^ in injury-induced ADM is sufficient to drive the phenotypic changes associated with PanIN formation, as well as the generation of MUC5AC+ pit-like cells. These data have important implications in regards to tumor cell of origin. Previous studies have focused on the role of acinar and ductal cells in tumorigenesis, however our data suggest that the plethora of cell types that form in response to injury are susceptible to oncogenic transformation. The particular cell state or cell type in which oncogenic mutation occurs may factor into the speed, aggressiveness, or sub-type of PDAC that develops. Further, these data shed new light into mechanisms by which pancreatitis predisposes patients to PDAC.

## Figure Legends

**Movie S1**. Animated visualization of the 3-D reconstruction of a metaplastic duct showing a heterogeneous consortium of cell types encapsulated by basement membrane with closely apposed vasculature. Volumetric reconstructions of cell plasma membranes and nuclei, as well as a portion of the basement membrane rendered in gray, demonstrate the morphological variability between cell types. Binucleate acinar cell, orange; mucinous cells, green; tuft cell, magenta; enteroendocrine cell, blue.

**Supplemental File 1:** Library QC Matrices

**Supplemental File 2:** DEGs_Injury vs. Normal

**Supplemental File 3:** EYFP-positive cell type markers

**Supplemental File 4:** EYFP-positive Regulons

**Supplemental File 5:** Gastric cell type signatures

**Supplemental File 6:** SPEM signatures

**Supplemental File 7:** Tuft and EEC markers

## Acknowledgments

We thank Richard DiPaolo, Christopher Wright, Eunyoung Choi, Ela Contreras Panta, and members of the Goldenring, Choi, and DelGiorno laboratories for helpful discussions. We thank Dr. Tom Bartol for his help with 3DEM stack alignment using SWiFT-IR (funded by NIH P41GM103712 and NSF DBI-1707356). Finally, we thank the scientists whose work this study builds on, but cannot be discussed here due to space constraints.

---

**Figure S1.**
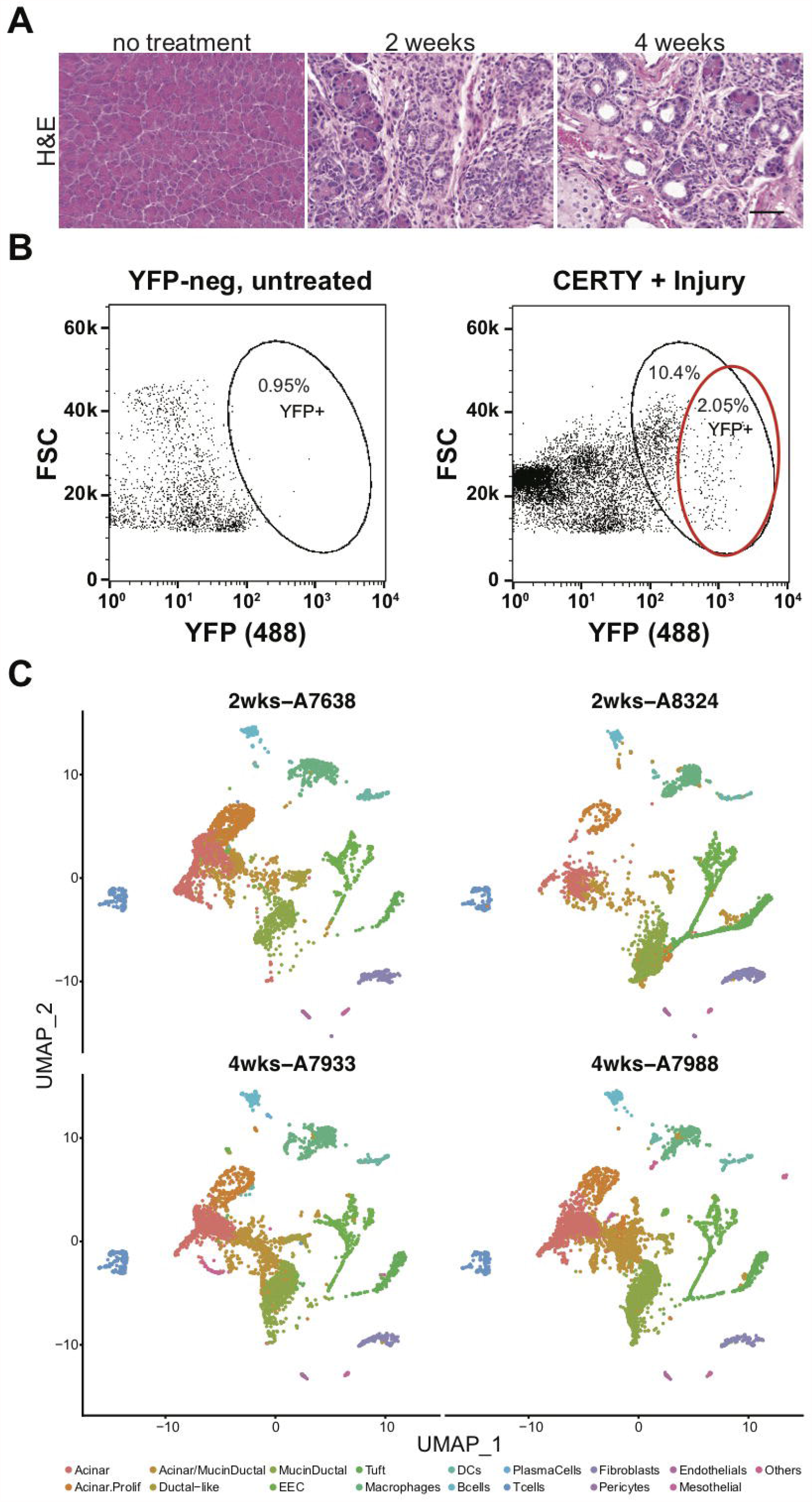
Isolation and analysis of lineage-traced EYFP+ cells from injured pancreata. **(A)** H&E analysis of the normal mouse pancreas or pancreata treated with caerulein for either 2 or 4 weeks. Scale bar, 50 µm. **(B)** FACS strategy for isolation of lineage traced EYFP+ cells from injured *CERTY* pancreata. Cells included in the red circle were collected to avoid injury-induced autofluorescence. **(C)** UMAPs showing annotated clusters from FACS collected EYFP+ cells from the injured pancreata of 4 *CERTY* mice treated with caerulein for either 2 or 4 weeks.

**Figure S2.**
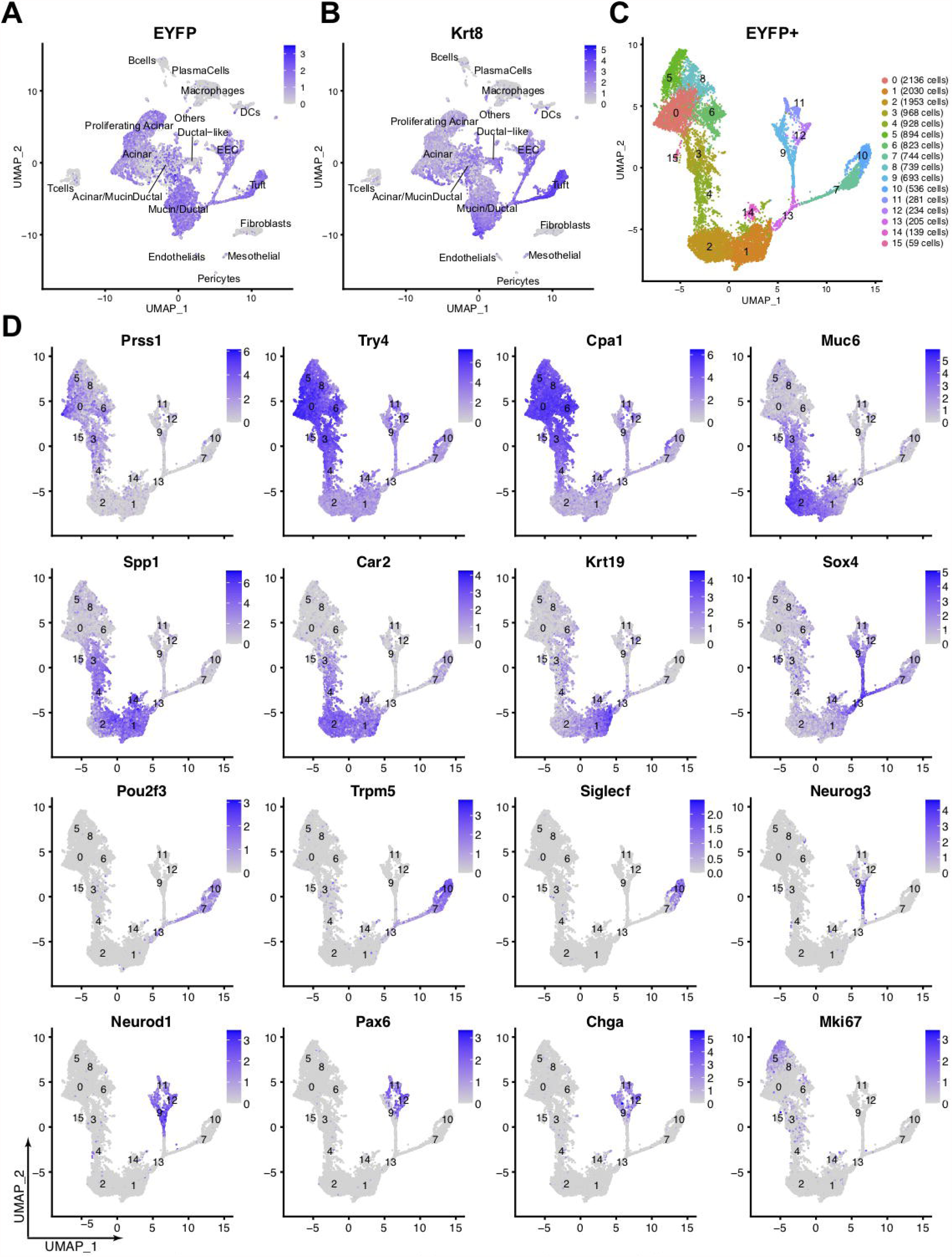
Annotation of lineage-traced EYFP+ cells. **(A)** Expression of *EYFP* and **(B)** epithelial cell marker *KrtB* overlaid on the UMAP of all FACS collected EYFP+ cells. **(C)** UMAP of EYFP+ cells from Figure 1B annotated with Seurat-identified clusters. **(D)** Expression of acinar cell *(Prss1, Try4* and *Cpa* 1), mucin/ductal cell *(Muc6, Spp1, Car2*, and *Krt19)*, tuft/EEC progenitor cell (*Sox4)*, tuft cell *(Pou2f3, TrpmS*, and *Siglecf)*, EEC *(Neurog3, Neuro d 1, Pax6*, and *Chga)*, and proliferation marker *Mki67* on the UMAP of EYFP+ cells shown in (C). Color intensity in A-C indicates the normalized gene expression level for a given gene in each cell.

**Figure S3.**
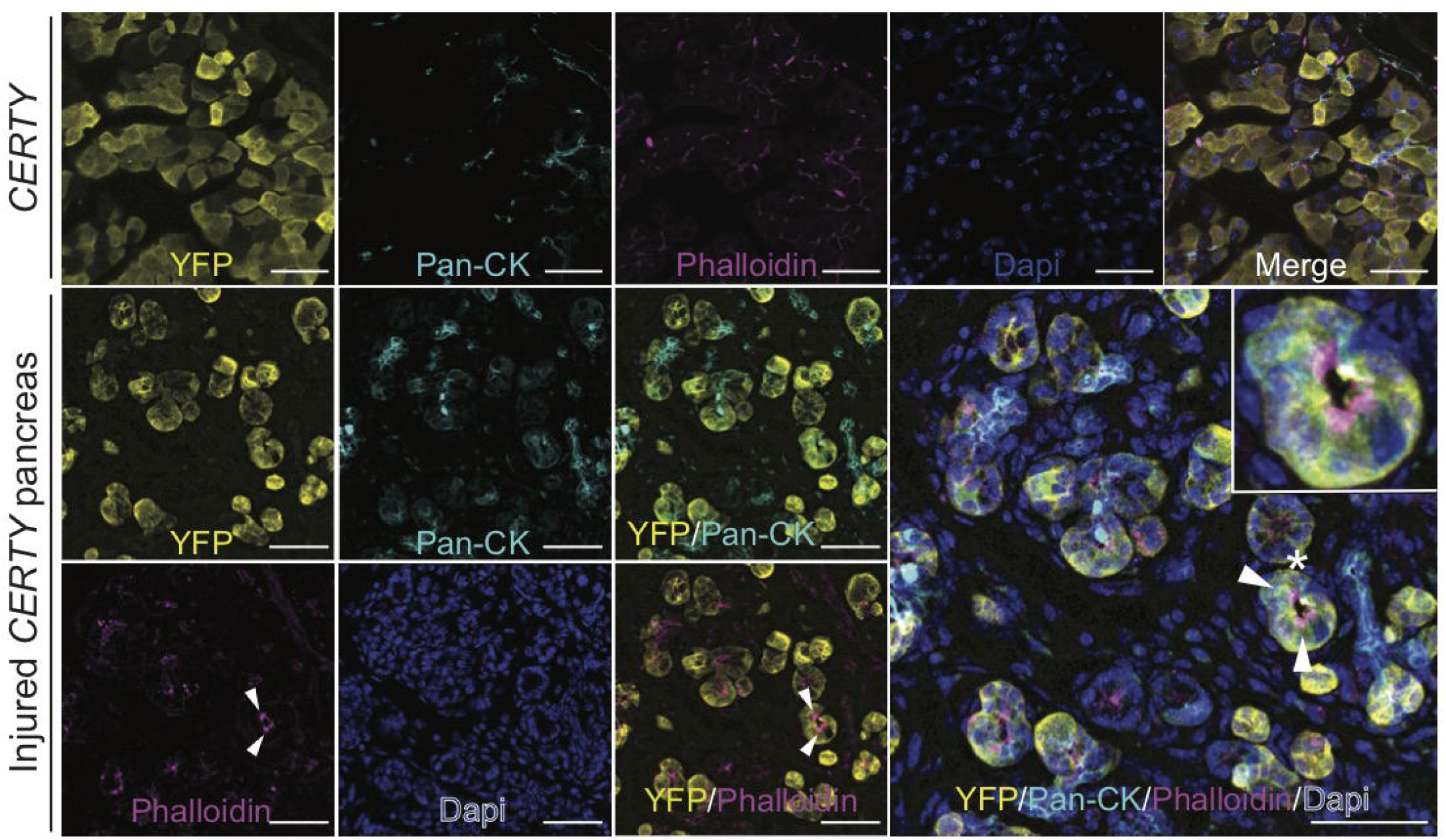
Acinar cell lineage tracing in normal and injured *CERTY pancreata*. Co-IF for EYFP (yellow), ductal cell marker PanCK (cyan), and membrane and tuft cell marker phalloidin (magenta) in normal or injured *CERTY* pancreata. Tuft cells highlighted in insert. Dapi, blue. Scale bars 100 µm.

**Figure S4.**
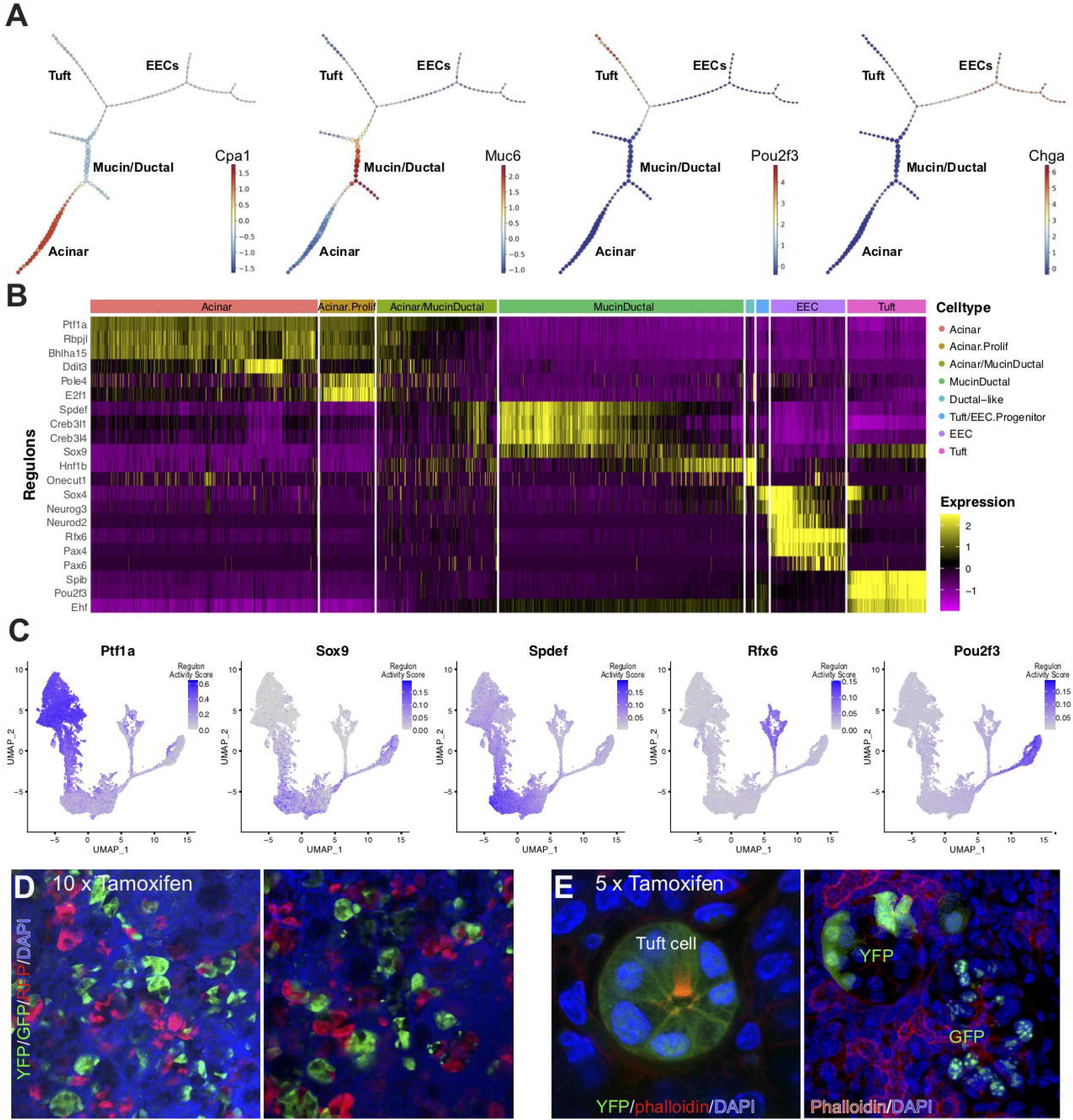
Lineage progression of ADM-derived populations. **(A)** Expression of acinar cell marker *Cpa1*, mucin/ductal marker *Muc6*, tuft cell marker *Pou2f3*, and EEC marker *Chga* overlaid on the pCreode plot shown in Figure 2C. **(B)** Heat map with hierarchical clustering showing regulons enriched in EYFP+ populations. **(C)** Expression of key transcription factors overlaid on the UMAP from Figure 1B. Color intensity indicates the relative enrichment of signature gene sets in each cell. **(D)** Expression of Brainbow reporters (YFP, green, cytoplasmic; GFP, green, nuclear; RFP, red, cytoplasmic) in injured pancreata from *Ptf1aCre*^*ERTM*^*;Rosa*^*Brainbow.2/+*^ mice treated with 10 or **(E)** 5 doses of tamoxifen. Phalloidin staining in (E) highlights a tuft cell. Dapi, blue.

**Figure S5.**
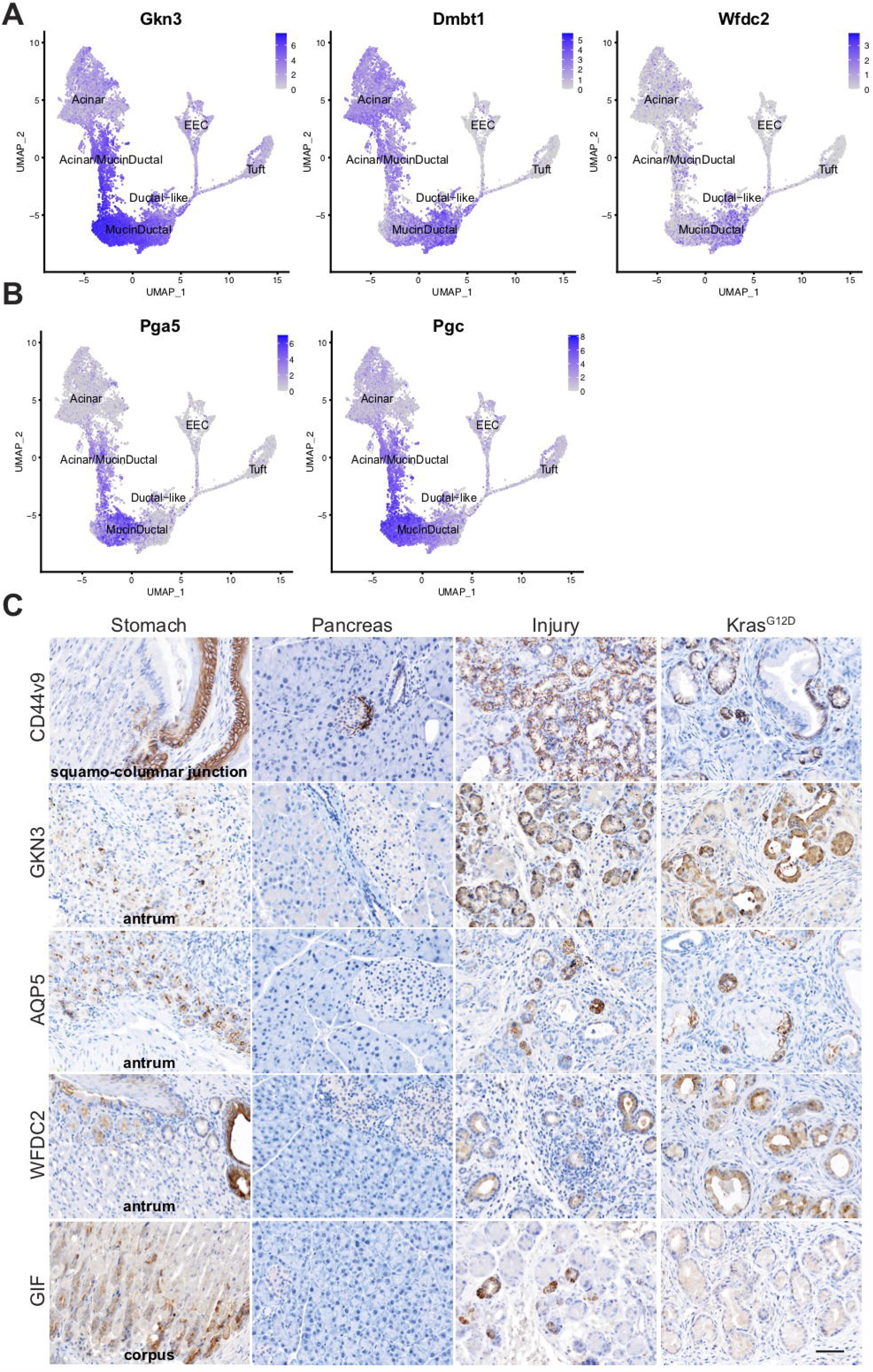
ADM as a pyloric-type metaplasia. **(A)** Expression of spasmolytic polypeptide-expressing metaplasia (SPEM) markers *Gkn3, Dmbt1*, and *Wfdc2* and **(B)** chief cell markers *Pga5* and *Pgc* overlaid on the UMAP of EYFP+ cells from Figure 1B. Color intensity indicates the relative enrichment of signature gene sets in each cell. **(C)** IHC for SPEM markers CD44v9, GKN3, AQP5, and WFDC2, and chief cell marker GIF in the normal stomach or pancreas and in injury-or oncogene-*(Kras*^*G12D*^ ) induced ADM and PanIN. I, islet. Scale bar, 50 µm.

**Figure S6.**
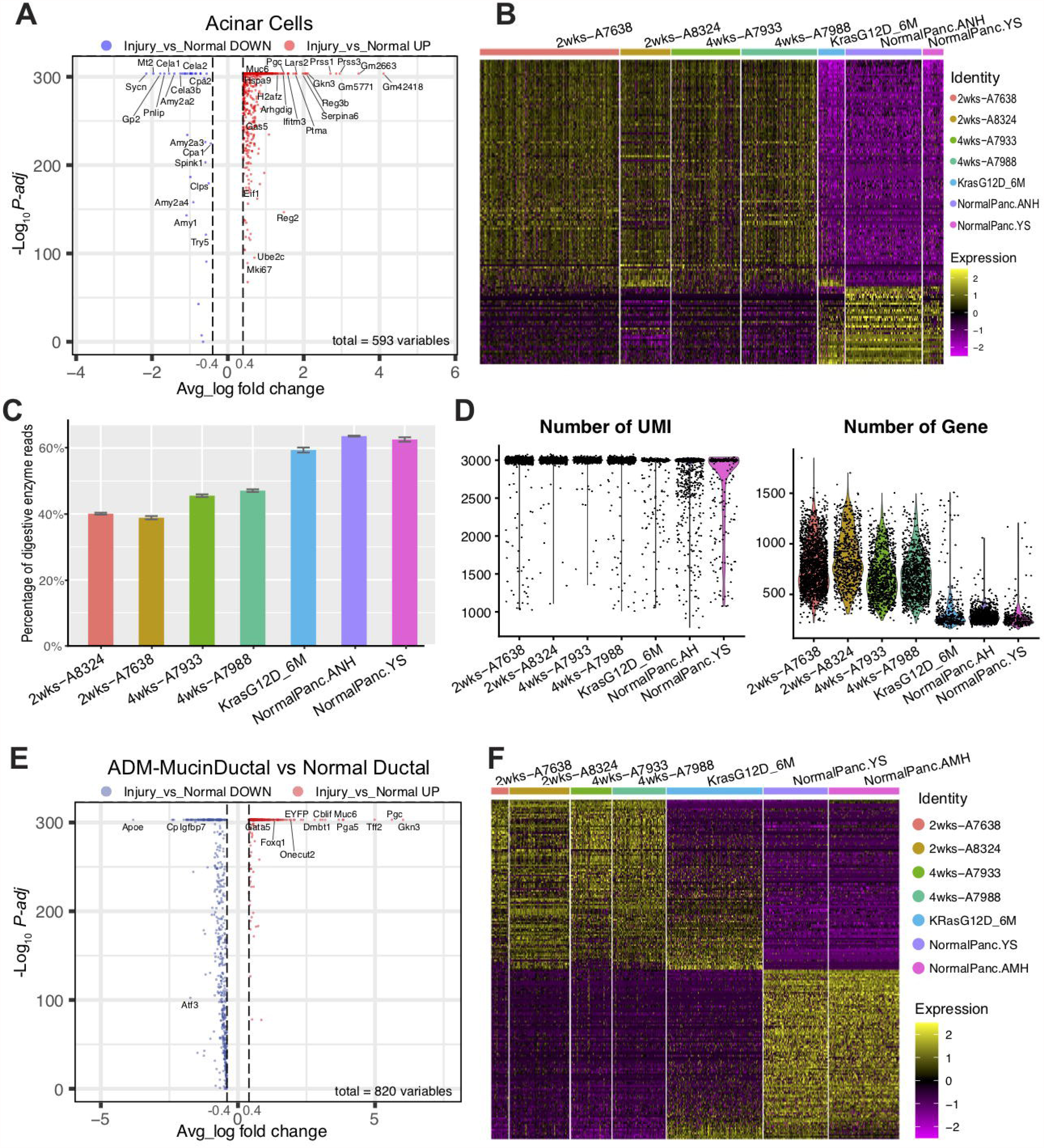
Comparison of normal and ADM-associated acinar and ductal cells. **(A)** Volcanoplot of the differentially expressed genes between injury acinar cells and normal acinar cells(Schesigner *et al*. (NormalPanc.YS) and Hosein *et* al.(NormalPanc.ANH)). **(B)** Heatmap of the top 100 up-regulated and all 34 down-regulated genes in A. **(C)** Barplot of the average percentage of transcripts from digestive enzymes in individual cells. Error bar represent the standard error. **(D)** Voilin plots of the number of UMI per cell and number of genes detected per cell across down-sampled (max nUMl=3k) samples. **(E)** Volcano plot of the differentially expressed genes between YFP+ ADM-induced mucin/ductal cells and normal ductal cells(Hendley *et al*., NormalPanc.AMH and Schlesigner *et al*., NormalPanc.YS). **(F)** Heatmap of the top 200 differentially expressed genes in E.

**Figure S7.**
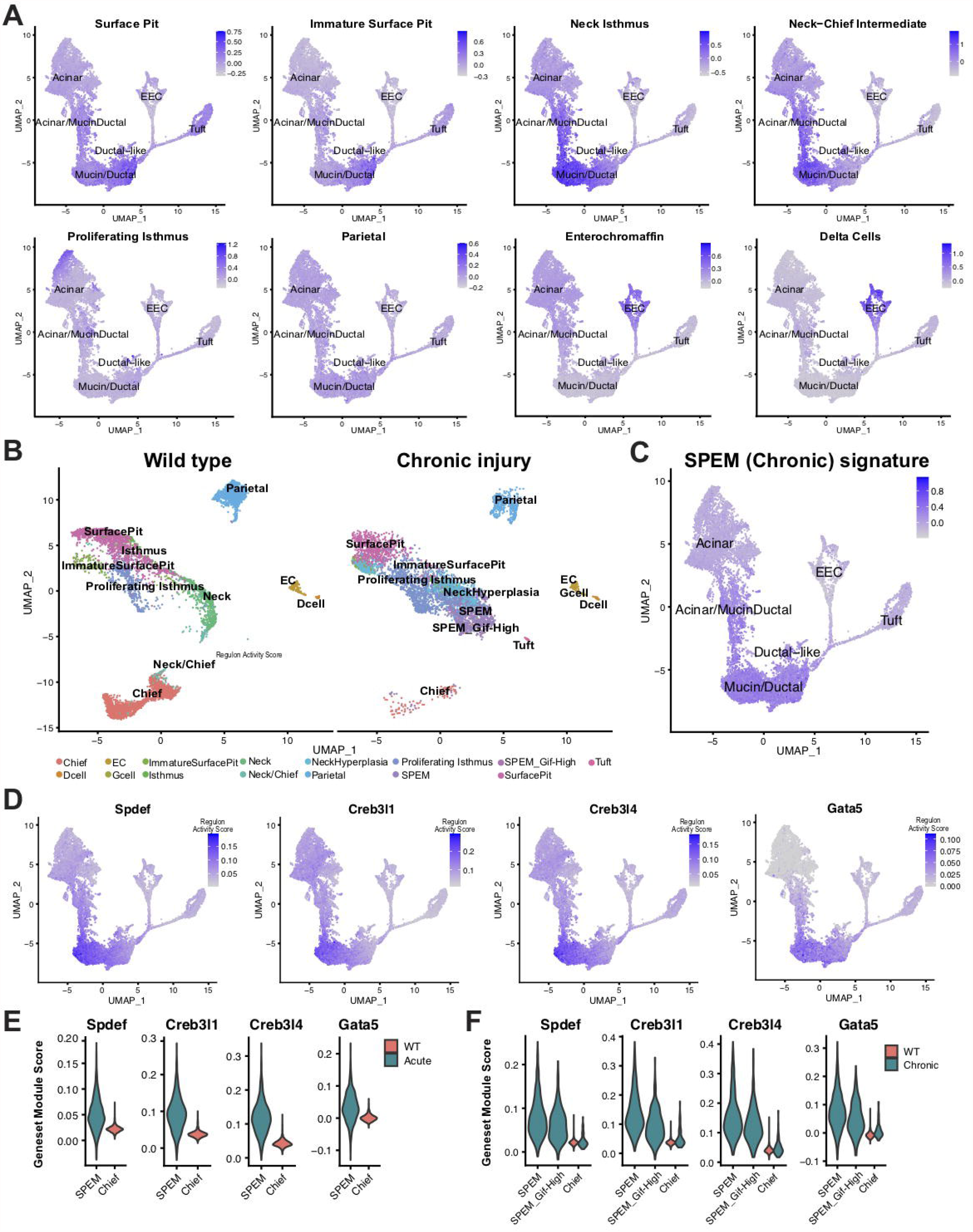
Comparison of ADM to normal and injured gastric epithelium. **(A)** Expression of normal gastric epithelial cell type signatures derived from Bockerstett et al., and shown in Figure 30 overlaid on the UMAP of EYFP+ cells from Figure 18. **(B)** UMAP of scRNA-seq data collected from normal stomach and a SPEM forming model of chronic gastric injury, derived from Bockerstett et al. **(C)** Expression of the chronic injury-induced gastric SPEM signature overlaid on the UMAP of EYFP+ cells from Figure 1B. **(D)** Regulon scores (transcription factor activity) overlaid on the UMAP from Figure 1B. **(E)** Gene module scores for TFs shown in (D) for normal gastric chief cells, acute SPEM cells, or **(F)** SPEM populations associated with chronic gastric injury. Color intensity in (A), (C), and (D) indicates the normalized gene expression level for a given gene in each cell. EC, enterochromaffin cell.

**Figure S8.**
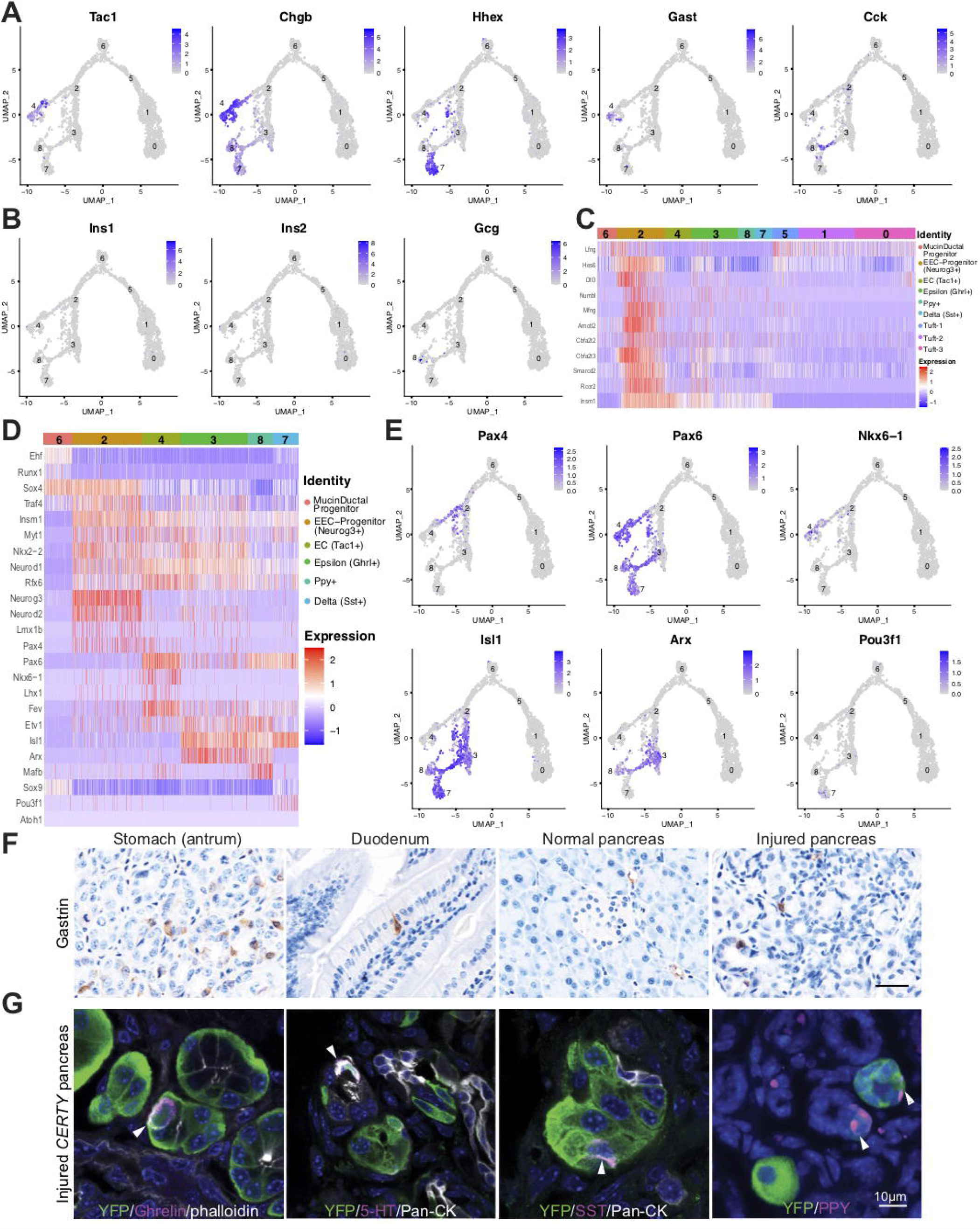
ADM results in substantial enteroendocrine cell heterogeneity. **(A)** Expression of EEC markers overlaid on the UMAP of tuft cells and EECs from Figure 4A. **(B)** Expressionof islet-derived hormones insulin *(lns1* and *lns2)* and glucagon (*Gcg*) overlaid on the UMAP from Figure 4A. **(C)** Heat map with hierarchical clustering showing expression of Notch pathway inhibitor genes enriched in EEC progenitor cells. **(D)** Heat map with hierarchical clustering showing key TFs expressed in Tuft/EEC progenitors, tuft cells, and EECs. **(E)** Expression of TFs enriched in different EEC populations, overlaid on the UMAP from Figure 4A. Color intensity indicates the normalized gene expression level for a given gene in each cell. **(F)** IHC for gastrin in the antrum of the normal stomach, duodenum, or pancreas and in the injured pancreas. Scale bar, 25 µm. **(G)** Lineage tracing and Co-IF for EYFP, green, hormones (Ghrelin, 5-HT, SST, PPY, magenta), pan-CK or phalloidin (white), or Dapi, blue. Scale bar, 10 µm.

**Figure S9.**
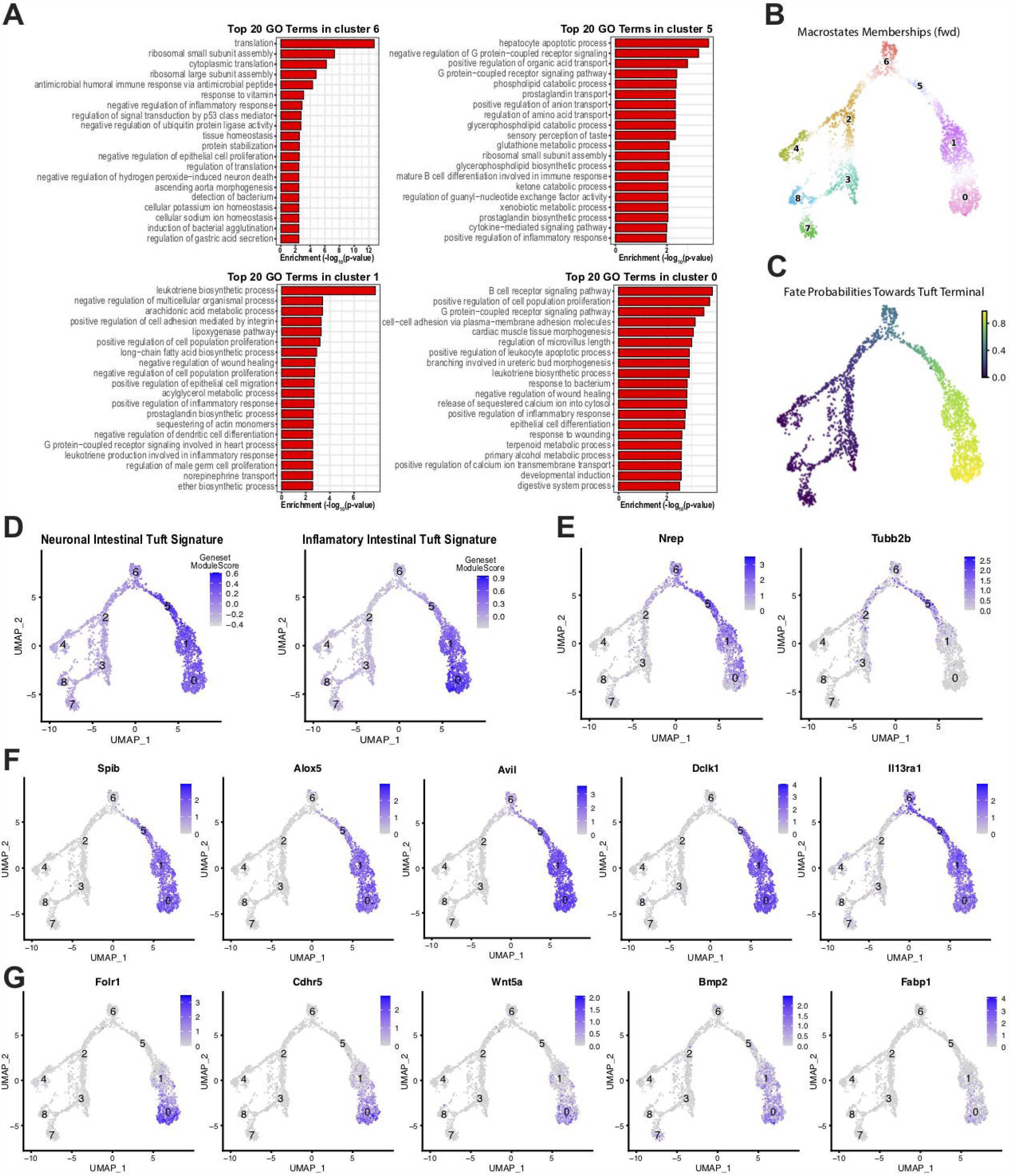
Tuft cell heterogeneity as a reflection of differentiation status. **(A)** Barplots of the enrichments scores of the top 20 GO terms associated with the DEGs in the 4 tuft cell clusters. The x-axis represents the enrichment score calculated by −log10(p-value of Fisher’s exact test). **(B)** 9 macrostates computed from Schur decomposition. The Seurat cluster labels were added to the associated macrostates and colored according to the Seurat cluster colors in Figure 4A. **(C)** Fate probability scores towards the terminal tuft cell lineage overlaid on the Tuft+EEC UMAP. **(D)** Seurat geneset module scores of the neuronal and inflammatory signatures in murine small intestinal tuft cells overlaid on the Tuft+EEC UMAP. **(E-G)** Expression of select genes overlaid on the Tuft+EEC UMAP, labeled by Seurat cluster.

**Figure S10.**
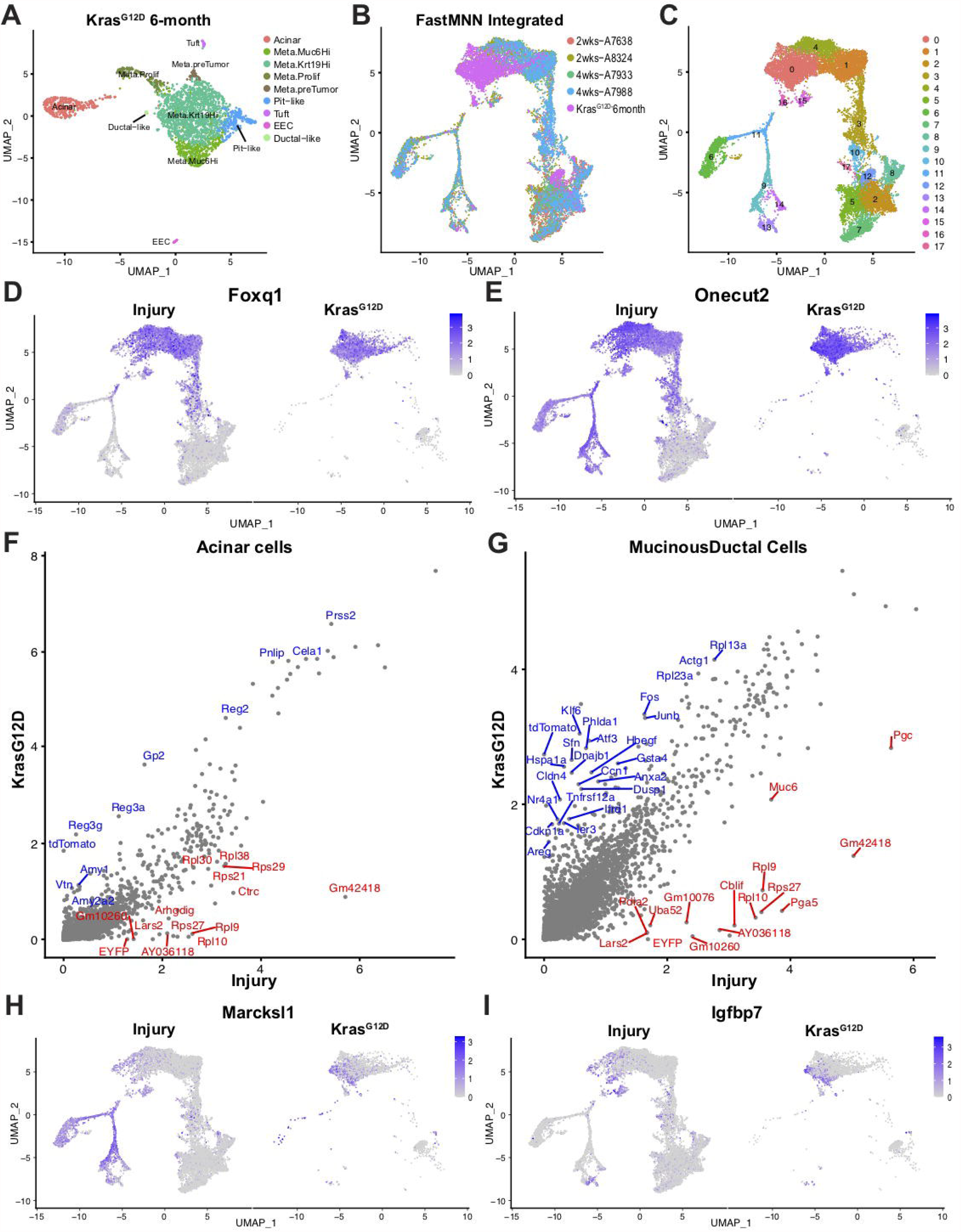
Comparison of injury and oncogene-induced ADM. **(A)** UMAP showing annotated clusters of scRNA-seq from the pancreas of a 6-month-old *KCT* mouse. **(B)** FastMNN integrated EYFP+ injury-induced ADM and tdTomato+ cells from a six-month-old *KCT* mouse, annotated by sample or **(C)** Seurat cluster. **(D)** Expression of metaplasia-specific transcription factors *Foxq1* and **(E)** *Onecut2* overlaid on the UMAP from Figure 6A. **(F)** Volcano plots showing the top 20 differentially expressed genes between the injury and oncogene-induced acinar and **(G)** mucin/ductal clusters. **(H)** Expression of pre-tumor markers *Marcks/1* and (I) */gfbp7* overlaid on the UMAP from Figure 6A. Color intensity indicates the normalized gene expression level for a given gene in each cell.

**Figure S11.**
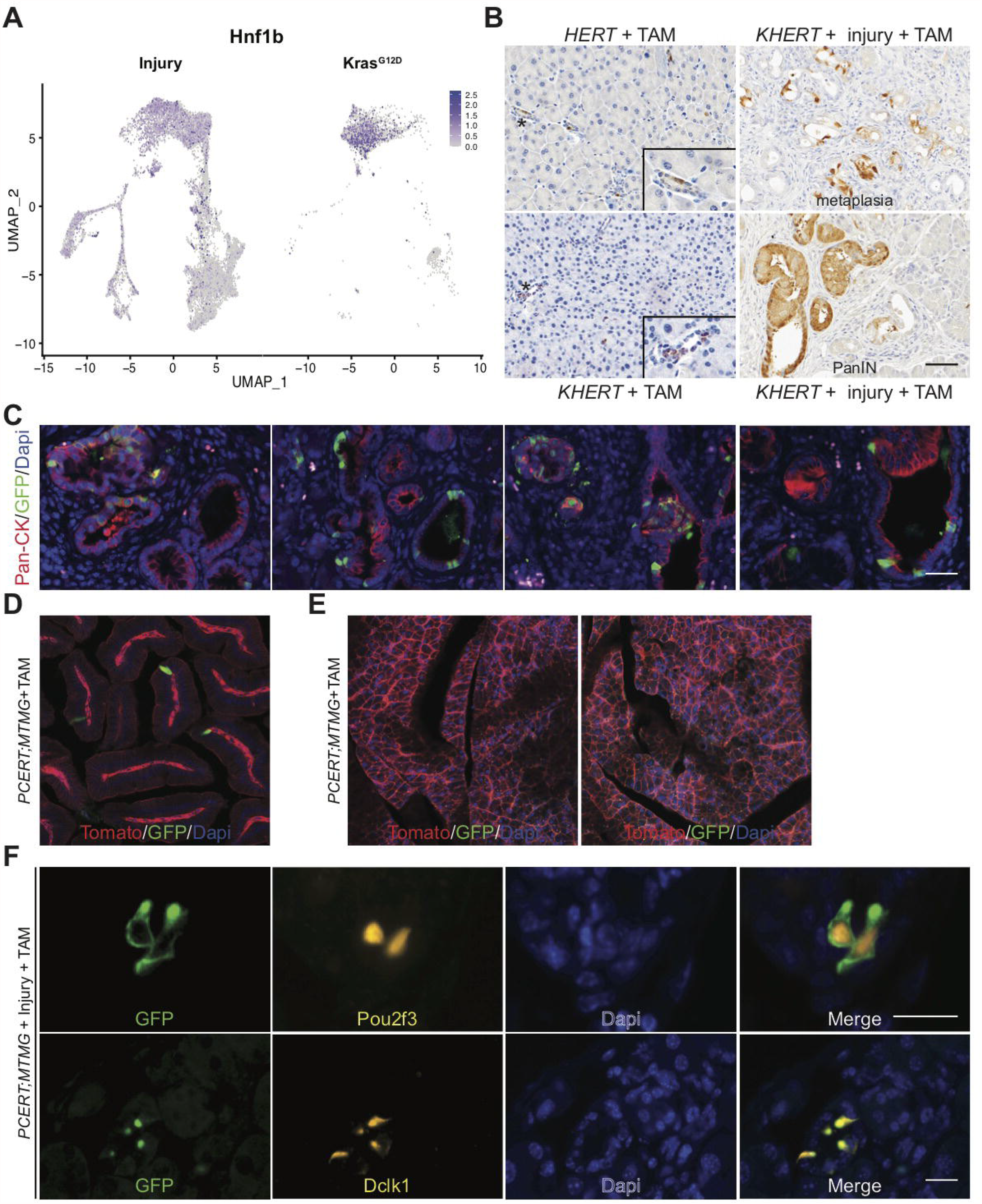
KrasG12D induction in ADM populations using inducible Cre mice. **(A)** Expression of *Hnf1b* overlaid on the UMAP from Figure 6A. **(B)** Lineage tracing and EYFP expression in *HERT* or *KHERT* mice treated with tamoxifen or *KHERT* mice first treated with caerulein to induce injury, followed by tamoxifen to induce *Kras*^*G12D*^ expression in ADM. Scale bar, 50µm. **(C)** Co-IF for EYFP (green) and ductal cell marker cytokeratin (pan-CK, red) in the pancreata of *KHERT* mice treated with caerulein then tamoxifen. Scale bar, 50µm. **(D)** Tomato (red) and GFP (green) expression in the intestines and **(E)** pancreas of a *PCERT;MTM G* mouse treated with tamoxifen. **(F)** Co-IF for GFP (green) and tuft cell makers POU2F3 or DCLK1 (both yellow) in the pancreas of a *PCERT;MTMG* mouse treated with caerulein to induce ADM and tuft cell formation, followed by tamoxifen to induce GFP expression in *Pou2f3+* cells. Scale bars, 25µm

**Figure S12.**
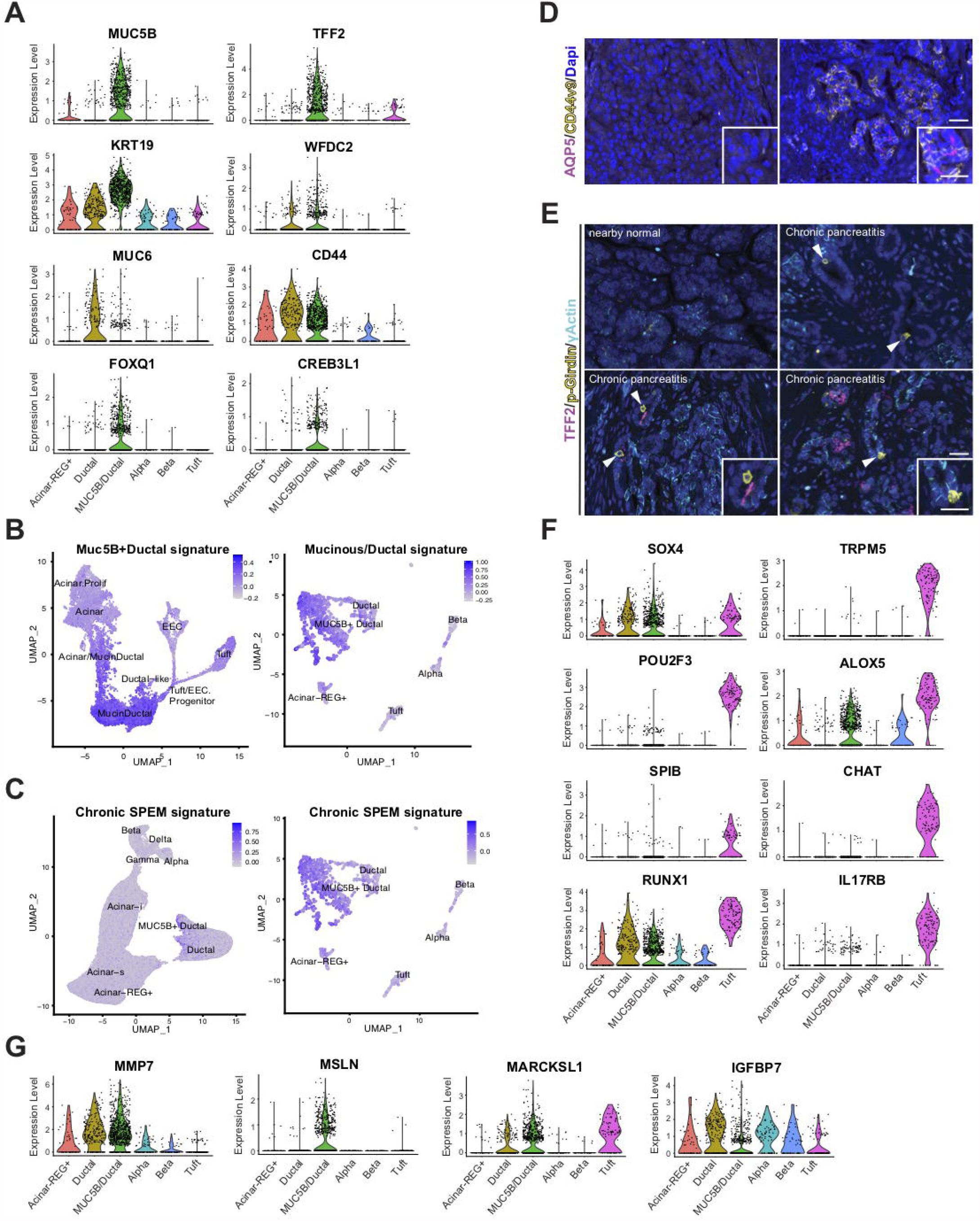
ADM cell type markers in Human pancreatitis. **(A)** Violin plots of MUC5B+Ductal cell markers *MUC58* and *KRT19*, SPEM markers *MUC6, TFF2, WFDC2*, and *CD44*, and mucin/ductal transcription factors *FOXQ1* and *CREB3L1* in sNuc-seq data from human pancreatitis. **(B)** Expression of the human MUC5B+Ductal signature overlaid on the mouse ADM dataset (left) and the murine mucin/ductal gene signature overlaid on the human chronic pancreatitis dataset (right). **(C)** Expression of the chronic SPEM signature shown in Figure S7 humanized and overlaid on sNuc-seq data from normal pancreas (left) and chronic pancreatitis (right). Color intensity indicates the relative gene expression for a given cell type in each cluster. **(D)** Co-IF for SPEM markers AQP5 (magenta), and CD44v9 (cyan), in an area of relatively normal pancreas (left) and injured pancreas (right) from the same patient sample. Scale bar, 25µm. **(E)** Co-IF for SPEM marker TFF2 (magenta), tuft cell marker phospho-Girdin (yellow), and y-actin in tissue specimens from patients with chronic pancreatitis. Scale bars, 25µm. **(F)** Violin plots of tuft cell markers and TFs *(SO X4, POU2F3, SP/8, RUNX1)* shown in Figure 5, or **(G)** pre-tumor markers in sNuc-seq data from human pancreatitis.

