## Supplemental Methods for "Single-cell transcriptomics reveals a conserved metaplasia program in pancreatic injury"

**Materials and Methods.**

**Mice.** Mice were housed in accordance with NIH guidelines in AAALAC-accredited facilities at Vanderbilt University, the Salk Institute, and at Washington University. The IACUC committees at Vanderbilt University, the Salk Institute, and Washington University approved all mouse procedures. *Ptf1a^Cre-ERTM^*, *Rosa^YFP/+^*, *Rosa^Brainbow.2/+^*, *Rosa^mT/mG^*, and *Hnf1b:CreER^T2^* mice were purchased from Jackson Laboratories^1, 2^. Mice were backcrossed into the CD-1 strain (Charles River). *Pou2f3-cre^Ert2^* mice have been reported previously, were provided by Dr. Ichiro Matsumoto (Monell Chemical Senses Center), and were backcrossed into the CD-1 strain^3^. For stomach studies, wild type C57BL/6 mice were purchased from The Jackson Laboratory and maintained in a specified-pathogen-free barrier facility under a 12-hour light cycle.

**Human samples**. The Institutional Review Board of Vanderbilt University approved the distribution and use of all human samples.

**Lineage tracing and pancreatitis induction.** Lineage tracing was conducted using the mouse model *Ptf1a^Cre-ERTM^*;*ROSA^YFP/+^*, as previously described^1, 4, 5^. In this model, tamoxifen treatment induces Cre activity, which then initiates expression of enhanced yellow fluorescent protein (EYFP) specifically in Ptf1a+ acinar cells. Acinar cells were labeled with daily tamoxifen treatment (Sigma, 5 mg/day, 5 days/week for two weeks) delivered in corn oil (Sigma) by oral gavage. Pancreatitis was induced with caerulein (Bachem) administered intraperitoneally (IP) at 250 μg/kg, twice a day, for five days a week for either two or four weeks and mice were then allowed to recover for two days^5^. Lineage tracing was conducted in *Ptf1a^Cre-ERTM^*;*ROSA^Brainbow.2/+^*mice with two daily treatments of 5 mg tamoxifen followed by the caerulein treatment protocol described above. Pancreatitis was induced in *Kras^G12D^*;*Hnf1b:CreER^T2^* mice (with or without *ROSA^YFP/+^*), *Hnf1b:CreER^T2^;Rosa^YFP/+^* mice, *Kras^G12D^*;*Pou2f3-cre^Ert2^* mice, or *Pou2f3-cre^Ert2^*;*Rosa^mTmG^* mice with caerulein administered at 250 μg/kg, twice a day, for five days a week for four weeks followed by tamoxifen treatment (5 mg/day, 5 days/week or tamoxifen chow for 5 days). Mice were allowed to recover for a minimum of 1 month or until they had to be sacrificed due to lung tumors (*Kras^G12D^*;*Hnf1b:CreER^T2^* mice) or large oral papillomas (*Kras^G12D^*;*Pou2f3-cre^Ert2^* mice).

**Isolation of pancreatic single cell suspensions.** EYFP+ cells were isolated from the pancreata of caerulein-treated mice. The pancreas was quickly dissected and minced in HBSS. Supernatant and fat were removed and pancreatic tissue was then incubated in 10 ml DMEM supplemented with 1 mg/ml collagenase I (Sigma), 1 mg/ml soybean trypsin inhibitor (Gibco), 0.5-1 mg/ml hyaluronidase (Sigma), and 250 μl of DNAse I, shaking gently at 37°C for a maximum of 30 min. Digestion was monitored and tissue was further digested mechanically by pipetting. Digested tissue was passed through a 100 μm filter, washed with PBS containing 1 mM EDTA and 0.5% BSA, and incubated with ACK lysing buffer (Gibco) to remove red blood cells. Single cell suspensions were incubated on ice with DAPI (molecular probes, 1:1000), used to exclude dead cells. EYFP+ cells were FACS purified at the Salk Institute’s Flow Cytometry core facility on a BD Biosciences Influx and an Aria Fusion cell sorter (100-µm size nozzle, 1 x PBS sheath buffer with sheath pressure set to 20 PSI). Cells were sorted in 1-drop Single Cell sort mode for counting accuracy.

**Single cell preparation and 10x 3’ library construction**. FACS-isolated EYFP+ cells were loaded onto a microfluidic chip with barcoded beads according to 10X Chromium Next GEM Single Cell 3’ (v3.1, Catalog # 1000128, 10x Genomics Inc.). Subsequent cell lysis, first strand cDNA synthesis and amplification were carried out according to the 10X v3.1 protocol, with cDNA amplification set for 11 cycles. Following sample indexing and bead-based library purification, the final library size distribution and concentration were measured using TapeStation (Agilent Biosystems) and Qubit (ThermoFisher). The libraries were pooled in equal molar ratio and sequenced on Illumina NovaSeq 6000.

**10X single cell data processing.** The raw sequencing data from each mouse was first processed separately in Cell Ranger 4.0.0 using a customized reference based on refdata-gex-mm10-2020-A-R26 to allow quantification of EYFP expression. The EYFP reporter transgene was added to the refdata-gex-mm10-2020-A-R26 reference and rebuilt by running cellranger mkref with default parameters (10x Genomics). Each dataset captured similar numbers of cells (~9k cells/mouse) with comparable sequencing depth and quality (see library QC summary in supplemental table1). The filtered gene-by-cell count matrices from 10x Cellranger were further QCed and analyzed in R package Seurat (v3.2.3)^6^. Each library was processed separately in Seurat with NormalizeData(normalization.method = "LogNormalize") function. The normalized data were further linear transformed by ScaleData() function prior to dimension reduction. Principal components analysis (PCA) was performed on the scaled data by only using the most variable 2000 genes (identified using the default “vst” method). Cells were examined across all clusters to determine the low quality cell threshold that accommodates the variation between celltypes. Low quality cells were removed with the same filtering parameters across libraries (percent.mt <=6 & nCount_RNA >= 1000). QCed dataset were reprocessed as described above and clusters were labeled with cell types based on marker gene expression and previously published scRNAseq derived gene signatures in pancreas. Doublets were marked using the doubetFinder R package (largely consistent with the results from Scrublet package) and removed. In addition to the doublets labeled by douletFinder. Two small clusters of cells with a high percentage of inferred doublets label, high number of UMI counts, and shared expression of marker genes were also removed as doublets. QCed datasets were integrated by using the fastMNN algorism included in the SeuratWrapper package to minimize the technical batch variations^7^. A union of the most variable 1500 genes from each sample was used as integration features for fastMNN. The first 45 MNN principal components were used in UMAP embedding and cluster identification. New clusters were re-labeled by checking both known individual marker gene expression and the module scores of previously generated cell-type-specific gene signatures from a pancreas scRNA-seq study^8^. EYFP negative clusters of stromal contamination were removed and the epithelium population was further filtered with EYFP transcripts resulting in 13362 EYFP+ cells (detectable EYFP transcripts) for downstream analysis.

Tuft and enteroendocrine cells were subsequently subset, re-scaled, and re-clustered to better reveal cellular heterogeneity. A small cluster of 107 cells become evident as doublets after re-clustering and was removed from subsequent analysis. This cluster expresses tuft, acinar, and mucin/Ductal cells and has a higher number of genes and UMI counts detected per “cell”. Differentially expressed genes were identified by using FindAllMarkers() function with logfc.threshold=0.5 and min.pct=0.2.

Processed human pancreas sNuc-seq dataset in seurat format from Tosti et al. was obtained from <http://singlecell.charite.de/pancreas/>. Gene signature enrichment was assessed by seurat geneset module score.

**GO enrichment analysis.** GO analysis was done with R package topGO v2.42.0 (algorithm = "elim", statistic = "fisher"))( <https://bioconductor.org/packages/release/bioc/html/topGO.html>). Genes meeting the filtering threshold in DEGs analysis (pct.min=0.2, min.diff.pct=-inf, min.cell.feature =3, logfc.threshold > 0) were used as allGenes to create the topGOdata object. The top 20 enriched GO terms were plotted in ggplot2. Enrichment score was calculated by taking -log10(p-value of Fisher’s exact test). In generating heatmaps of selected genes, cells were grouped by clusters and cells within a cluster group were re-ordered by increasing pseudotime within each cluster to better reveal expression pattern changes.

**RNA velocity estimation.** RNA velocity estimation was performed separately in each sample using the Velocyto.R package (v0.6)^9^. The spliced/unspliced/ambiguous reads classification was done by running the velocyto run10x pipeline on cellranger output. The resulting loom file was converted to Seurat object and subset to keep the 13362 EYFP+ cells. RNA velocity estimation was performed by using RunVelocity() function in Velocyto.R package with the following parameters: deltaT=1, ambiguous="ambiguous", kCells=25, fit.quantile=0.02,reduction = "mnn", ncores=30. Cell-cell distance used in the Runvelocity() step was calculated from the fastMNN corrected principle components space. Calculated velocity was visualized on the UMAP embedding through the show.velocity.on.embedding.cor() function with the parameters setting: min.grid.cell.mass=0.5, show.grid.flow=TRUE, grid.n=40, do.par=T, n.cores=30,cell.border.alpha=0.1, arrow.scale=1.

**CellRank analysis**

The aggregate fate probabilities of tuft+eec lineage were calculated using the CellRank package (v.1.2.0) (**doi:** https://doi.org/10.1101/2020.10.19.345983). Briefly, the count matrix from the RNA assay in the Seurat object of Tuft+EEC data subset was converted into h5ad format using the SeuratDisk package and subsequently loaded using scranpy for CellRank analysis. New neighborhood graph was recomputed in scranpy using the first 30 fastMNN principal components. To get cell-cell transition probabilities, we used a PalantirKenel on monocle3 psudotime. 9 macrostates corresponding to the 9 seurat clusters were identified using the compute_macrostate() function in the GPCCA estimator of the CellRank package. Macrostates corresponding to the Seurat clusters 0, 4, 7, and 8 were labeled as terminal states and the fate probabilities towards the tuft terminal state (seurat cluster 0) was calculated using the pl.cluster_fates()function. Similarly, the regulon activity score matrix was imported into scranpy for plotting regulon acvitity score over monole3 pseudotime. The plot was generated using the pl.gene.trends(n_test_point=500) function by fitting a GAM model between regulon activity score and monocle3 pseudotime in the cellrank package.

**Regulon Analysis**

Gene regulatory network (Regulon) analysis in this study was performed using the SCENIC R package (v1.2.0)^10^ The mouse mm10 500bp_up_100bp_down and 10kb_up_and_down databases were downloaded from the cistarget database; the TSS database was downloaded from the Aerts lab website (<https://resources.aertslab.org/cistarget/>) and used in this analysis. The filtered count matrix of the 13362 EYFP+ cells were further filtered to exclude non-expressed genes using the function geneFiltering(exprMat, scenicOptions=scenicOptions, minCountsPerGene=100, minSamples=25) in the SCENIC package. Spearman correlation between each TF and other genes was run by runCorrelation() function with the default settings. TF-target gene co-expression module was identified in pySCENIC by running the Run_arboreto_with_multiprocessing.py pipeline with default setting and random seed 915. The adjacency file was imported into SCENIC R package and the normalized weight was used as weight subsequent steps. Regulon and regulon activity was calculated by running the runSCENIC_1_coexNetwork2modules(), runSCENIC_2_createRegulons(),runSCENIC_3_scoreCells() functions with default settings. Filtered Count matrix was used in step 3 for regulon activity scoring using the AUCell algorism. The resulting regulon activity score matrix was subsequently imported into Seurat through CreateAssayObject() function. Regulon activity score was plotted in Seurat without further normalization or scaling.

**Monocole3 Trajectory analysis**

Seurat object of the 13362 EYFP+ cells was converted into Monocle3 CellDataSet object by using the as.cell_data_set()function in the SeuratWrapper Package. Subsequent trajectory analysis is done in the monocle3 package^11^. Cells were re-clustered in monocle by running the cluster_cells() function with setting resolution= 2.5e-4. Then cells were ordered in pseudotime by running the order_cells() function with default setting. Root state was manually set to acinar cells. Trajectory plot was generated using the plot_cells() function. Tuft+EEC monocle3 trajectories was done similarly as above. 2586 Tuft+EEC cells subset was used and the cluster_cells() function was run with resolution = 1e-3.The root state was manually set to Cluster 6 based on the RNA velocity information.

**p-Creode trajectory analysis.** Count matrices were first filtered using the dropkick algorithm to identify cells versus ambient barcodes^12^. Counts data were then normalized to the median total read count across all retained barcodes, transformed using an inverse hyperbolic sine function, and Z-score standardized through functions available in Scanpy (v1.6.1)^13^. p-Creode (v2.2.0) was performed on normalized data in supervised mode with a max of 10 principal components across 1000 scored bootstrapped runs. The highest scoring run, producing the most representative graph from the ensemble, was chosen to visualize overlays and pseudotime gene expression dynamics. These gene expression dynamics were smoothened and plotted using a generalized additive model, implemented through pyGAM (v0.8.0, zenodo.org). p-Creode density parameters were chosen according to automatic thresholding, or the best_guess function. Noise and target density parameters varied between the different datasets analyzed, resulting in the use of approximately 20% to 25% of cells in each bootstrapped replicate. Higher-resolution runs of p-Creode were performed on an enteroendocrine identity-enriched subset of cells by selecting cell populations with specific expression of tuft or enteroendocrine markers.

**Data integration of injury 10x scRNA-seq data and published datasets**

To avoid artifacts in differential gene analysis between our dataset and previous published datasets, fastq files were downloaded from the SRA using accession number GSE141007 (Schlesinger et al.), GSE159343 (Hendley et al.) and subsequently reprocessed using the same version of mm10 reference genome annotation (refdata-gex-2020-A reference genome from 10x Genomics) as our injury dataset. tdTomato transgene were added to the reference genome to allow quantification of tdTomato transgene. The Kras^G12D^ datasets were QCed using similar thresholds as that were used in the original study (minimum of 200 genes and 1000 UMIs per cell) except a slightly higher cutoff threshold for percent of mitochondrial reads (<=7.5%). Kras^G12D^ dataset and our Injury dataset were merged in seurat v3 and further integrated using the fastMNN method. For integration of our injury dataset and the sorted 6month Kras^G12D^ dataset, top 2000 variable genes identified using the “vst” method on the merged Injury+Kras^G12D^ dataset were used as the integration feature for fastMNN. The first 30 PCs returned by the fastMNN algorism were used for 2-dimensional UMAP embedding and Cluster identification. Normal pancreas scRNA-seq dataset from Hendley et al was QCed with percent.mt <=10 & nCount_RNA>=2000 & nCount_RNA < 30000. Cells were clustered and annotated using classic markers.

For comparison between injury and normal pancreas, population of interest were extracted into a separate seurat object and differential gene expression was performed on the merged injury+normalPancreas dataset between the injury group and normal group using FindMarkers() function in seurat with logfc.threshold=0.4. min.pct=0.3 was used for the ductal cell comparison and min.pct=0.2 was used for acinar cell comparison.

**Histological staining.** Tissues were fixed overnight in zinc-containing, neutral-buffered formalin (Fisher Scientific), embedded in paraffin, cut in 5 μm sections, mounted, and stained. Sections were deparaffinized in xylene, rehydrated in ethanol, and then washed in PBST and PBS. Endogenous peroxidase activity was blocked with a 1:50 solution of 30% H_2_O_2_:PBS followed by microwave antigen retrieval in 100 mM sodium citrate, pH 6.0. Sections were blocked with 1% bovine serum albumin (BSA) and 5% normal goat or rabbit serum in 10 mM Tris (pH 7.4), 100 mM MgCl_2_, and 0.5% Tween-20 for 1hr at room temperature. Primary antibodies were diluted in blocking solution and incubated overnight. Anti-Gkn3 was generated by the Mills Laboratory (Washington University, St. Louis, MO), anti-Tff2 was a generous gift from Sir Nicholas Wright (Barts Centre, London), and anti-Gif was a generous gift from the Alberts Laboratory (Washington University, St. Louis, MO). Information on additional primary antibodies is provided in Table S1. Slides were then washed, incubated in streptavidin-conjugated secondaries (for rabbit or mouse antibodies, Abcam, for rat or goat antibodies, Vector) and developed with DAB substrate (Vector). Hematoxylin and eosin (H&E) staining was performed to assess tissue morphology. All slides were scanned and imaged on an Olympus VS-120 or Olympus VS-200 Virtual Slide Scanning microscope.

**Fluorescence microscopy.** Tissues were fixed overnight in 4% paraformaldehyde, washed 3x with PBS and floated overnight in 30% sucrose. Tissues were then incubated in a 1:1 mixture of 30% sucrose and Tissue-Tek optimal cutting temperature compound (OCT, VWR) for 30 min, embedded in OCT and frozen at -80°C. 7 μm tissue sections were cut, permeabilized with 0.1% Triton X-100 in 10 mM PBS, and blocked with 5% normal donkey serum and 1% BSA in 10 mM PBS for 1 hour at room temperature. Tissue sections were stained with primary antibodies in 10 mM PBS supplemented with 1% BSA and 0.1% Triton X-100 overnight (Table S1). Sections were then washed 3 x 15 min in PBS with 1% Triton X-100, incubated in Alexa Fluor secondary antibodies and/or phalloidin (Invitrogen), washed again for 3 x 5 min, rinsed with distilled water, and mounted with Prolong Gold containing Dapi (Invitrogen). Immunofluorescence on paraffin-embedded tissues followed the immunohistochemistry protocol until the blocking step. Instead, tissues were blocked in the donkey serum block described above and then followed the protocol for fluorescence microscopy described here. All slides were imaged on an Olympus VS-120, an Olympus VS-200 Virtual Slide Scanning microscope, or Zeiss 880 Airyscan Super-Resolution microscope.

For multiplex staining featured in Figure 2B, primary antibodies (Supplemental Table 1) were used at the listed concentrations and incubated in tris buffered saline (TBS) overnight. The next day slides were washed 4 x 5 min in TBS and incubated for 1 hour at room temperature with DAPI (H-1000, Invitrogen) and secondary antibodies raised in donkey or goat and conjugated to AlexaFluor 488, 555, 594 or 647 (Invitrogen or JacksonImmunoresearch) were used at 1:400 dilution. After incubation, samples were washed 4x 5 min in TBS, covered with Vectashield (Invitrogen) and cover-slipped. Multi-plant confocal images were acquired using an up-right Leica Stellaris X5 microscope with a 20x/0.75 N.A. multi-immersion objective. After acquisition, images were processed using ImageJ/Fiji (NIH) and denoised using a median filter of 1px. Images shown are maximum projections of confocal stacks.

**instant Structured Illumination Microscopy (iSIM).** Image-stacks of Brainbow lineage tracing were captured with a system assembled by BioVision Technologies (Exton, PA) consisting of a VisiTech International (Sunderland, United Kingdom) iSIM module attached to a Nikon (Melville, NY) Ti2 stand equipped with a Nikon 100x 1.49 Plan Apo objective. Raw image stacks were then deconvolved using a Microvolution (Cupertino, CA) plugin in Fiji (ImageJ).

**3DEM Methods.** Samples of injured pancreas tissue were processed and imaged as described previously with some modifications^14^. Materials were sourced from Electron Microscopy Sciences (Hatfield, PA) unless noted otherwise. Briefly, fresh biopsies of injured tissues were dissected into small pieces (<1mm) and quickly immersed in 37°C buffered fixative (2.5% glutaraldehyde, 2% paraformaldehyde, 3mM CaCl_2_, 0.1M sodium cacodylate) before overnight storage at 4°C in the same solution^15^. Samples were serially rinsed with ice-cold cacodylate/CaCl_2_ buffer and stained with buffered reduced osmium (1.5% Osmium, 1.5% potassium ferrocyanide, 3mM CaCl_2_, 0.1M sodium cacodylate) for 45 minutes in the dark at room temperature. Samples were serially rinsed with ice cold distilled water before 30 minutes of treatment with filtered 1% aqueous thiocarbohydrazide (Ted Pella) that was prepared at 60°C for an hour before use. Samples were serially rinsed with ice cold distilled water before staining with 1.5% aqueous osmium tetroxide for 45 minutes in the dark at room temperature. Samples were rinsed serially with ice cold water before staining overnight with 1% uranyl acetate at 4°C. Samples were rinsed with distilled water and stained with Walton’s lead aspartate for 30 minutes at 60°C^16^. Samples were rinsed again with distilled water and serially dehydrated in ice cold ethanol. Samples were then infiltrated with Durcupan resin (hard formulation) and polymerized at 60°C for 3 days.

A sample block was prepared for serial sectioning as previously described and a ribbon of 248 serial sections, each of dimension ~150µmx400µmx70nm (x-y-z), were collected onto a silicon wafer diced to a dimension of 35x6mm using Diatome diamond knives mounted in a Leica UC8 ultramicrotome^17, 18^. The chip was imaged using a Zeiss Sigma VP scanning electron microscope equipped with a Gatan backscattered electron detector and Atlas5 (Fibics) control software. Multi-resolution maps were generated for a subset of sections throughout the ribbon, of which images were used for Figure 2, and a region of interest (ROI) for 3DEM was identified. Images were acquired with an accelerating voltage of 3kV, a 20µm aperture, at a working distance of 9mm, and with pixel size of 8nm. Contiguous images of the ROI were assembled into a series of 8-bit .tiff and aligned.

A delta cell, a tuft cell, two mucinous cells, an acinar cell, and their associated basement membrane were identified by their salient ultrastructural features. Their boundaries, along with their respective nuclei, were traced using VAST Lite annotation software^19^. Cells and nuclei were exported using the Matlab tools bundled with VAST Lite as .obj files, which were imported into Blender 2.79 (Blender Foundation) with add-ons BlendGAMer and Neuromorph both installed^20, 21^. Cells and nuclei were processed with GAMer to optimize geometry for visualization, before superimposition with 3DEM data using Neuromorph and animation and rendering using Blender. For video figures, rendered scenes were assembled into .avi files using ImageJ, post-processed using After Effects (Adobe), and compressed using Handbrake (Handbrake.fr)^22^.

**Transmission electron microscopy of murine pancreas**

Tissues were fixed in a solution of 2% paraformaldehyde, 2.5% glutaraldehyde, and 2 mM CaCl_2_ in 0.15 M sodium cacodylate buffer (pH 7.4) for 2 hours at room temperature. They were then post-fixed in 1% osmium tetroxide for 40 min and 1.5% potassium ferricyanide in sodium cacodylate buffer for 1 hr at 4°C in the dark. Tissues were stained en bloc in 1% aqueous uranyl acetate (4C in the dark), dehydrated in a series of graded ethanols, and embedded in Eponate12 resin (Ted Pella). Ultra-thin sections (70 nm) were obtained using a diamond knife (Diatome) in an ultramicrotome (Leica EM UC7) and placed on copper grids (300 mesh). Sections were imaged on a Zeiss Libra 120 TEM operated at 120 kV.

**Generation of SPEM and transmission electron microscopy of murine stomach.** Gastric SPEM was induced with tamoxifen (5mg/20g mouse body weight; Toronto Research Chemicals Inc., Toronto, Canada) administered by intraperitoneal injection for three consecutive days^23^. Tamoxifen was prepared by sonication in a 90% sunflower seed oil (Sigma, St Louis, MO) and 10% ethanol solution. For transmission electron microscopy, stomachs were prepared and imaged as previously described^24^. Briefly, stomach corpus was collected as described, fixed overnight at 4˚C in fixative (modified Karnovsky’s), and sectioned. Tissue was processed by the Washington University in St. Louis Department of Pathology and Immunology Electron Microscopy Facility.

All images (IHC, IF, EM) were digitally enhanced to edit the color, brightness and contrast levels using Zen (Carl Zeiss), ImageJ/Fiji (NIH), and/or Photoshop (Adobe) software.

**Figures.** Models shown in Figures 1A and in the graphical abstract were generated using BioRender.com.
