## Supplemental Table 1 for "Single-cell transcriptomics reveals a conserved metaplasia program in pancreatic injury"

| **Antibody** | **Company** | **Catalog #** | **Dilution, IF** | **Dilution, IHC** |
| --- | --- | --- | --- | --- |
| Amylase | Abcam | ab21156 | N/A | 1:5000 |
| Aqp5 | Sigma | HPA-065008 | 1:500 | 1:500 |
| Cd44v9 (anti-mouse) | Cosmo Bio Co. | LKG-M002 | 1:10,000 | 1:10,000 |
| Cd44v9 (anti-human) | Cosmo Bio Co. | LKG0M003 | 1:10,000 | N/A |
| ChgA | Santa Cruz | sc-1488 | 1:100 | N/A |
| Gamma actin | Santa Cruz | sc-65638 | 1:100 | N/A |
| Gastrin | Biogenex | PU019-UP | N/A | 1:100 |
| GFP | Abcam | ab6673 | N/A | 1:1000 |
| Ghrelin | Cell Signaling | 31865 | 1:1000 | 1:2000 |
| GIF | N/A | N/A | 1:1000 | 1:1000 |
| Gkn3 | N/A | N/A | N/A | 1:150 |
| MSLN (anti-mouse) | LS Bio | LS-C407883 | N/A | 1:2000 |
| Muc5ac | Thermofisher | MA5-12178 | N/A | 1:500 |
| Pan-CK | Abcam | ab9377 | 1:100 | N/A |
| Phospho-Girdin (pY1798) | IBL  America | 28143 | 1:100 | N/A |
| PPY | Abcam | ab255827 | 1:80,000 | 1:80,000 |
| RFP |  |  |  |  |
| Serotonin | Santa Cruz | sc-58031 | 1:200 | 1:500 |
| Somatostatin | Millipore | MAB354 | 1:1000 | 1:2000 |
| Synaptophysin | Cell Marque | 336R-94 | 1:500 |  |
| TFF2 | N/A | N/A | 1:500 | N/A |
| Wfdc2 | Invitrogen | PA5-80227 | N/A | 1:2000 |

Table S1. Primary antibodies used in immunofluorescence (IF) and immunohistochemistry (IHC) studies.
